# Metagenome-assembled microbial genomes from Parkinson’s disease fecal samples

**DOI:** 10.1101/2023.02.27.526590

**Authors:** Ilhan Cem Duru, Alexandre Lecomte, Tânia Keiko Shishido, Pia Laine, Joni Suppula, Lars Paulin, Filip Scheperjans, Pedro A. B. Pereira, Petri Auvinen

## Abstract

The human gut microbiome composition has been linked to Parkinson’s disease (PD). However, knowledge of the gut microbiota on the genome level is still limited. Here we performed deep metagenomic sequencing and binning to build metagenome-assembled genomes (MAG) from 136 human fecal microbiomes (68 PD samples and 68 control samples). We constructed 952 non-redundant high-quality MAGs and compared them between PD and control groups. Among these MAGs, there were 22 different versions of *Collinsella* and *Prevotella* MAGs, indicating high variability of those genera in the human gut environment. Microdiversity analysis indicated that *Ruminococcus bromii* was statistically significantly (p < 0.002) more diverse on the strain level in the control samples compared to the PD samples. In addition, by clustering all genes and performing presence-absence analysis between groups, we identified several control-specific (p < 0.05) related genes, such as *speF* and Fe-S oxidoreductase. We also report detailed annotation of MAGs, including Clusters of Orthologous Genes (COG), Cas operon type, antiviral gene, prophage, and secondary metabolites biosynthetic gene clusters, which can be useful for providing a reference for future studies.

## Introduction

Parkinson’s disease (PD) is a common neurodegenerative disease. Several studies have shown that gut microbiota have an impact on human health [^1,2^], as changes in the gut microbiota composition affect the brain–gut communication and pathology [^2^]. Accordingly, different neurodegenerative diseases, especially PD, are associated with gut microbiota composition changes [^3^]. Although several studies have shown which gut microbiota mechanisms are potentially relevant to PD [^3,4^], more comprehensive knowledge of the gut microbiota on the genome level could help to understand the functional impact of gut microbiota in PD.

With respect to gut microbiota, the Helsinki cohort is one of the most thoroughly studied cohorts in the field of PD. We have originally described differences between PD and control groups from stool samples [^3^], and followed the temporal stability of the microbiota [^4^]. In addition, the bowel symptoms and their linkage to the gut microbiota have been studied [^5^]. Following the Braak hypothesis, we have also investigated the microbiome and PD connections in samples from nasal and oral locations [^6^]. These original microbiome analyses have been followed by studies on SCFA and immune responses [^7^], and recently on the links between metabolomics, metabolic enzymes, and epigenetic regulation of the host [^8–10^].

In this study, we used shotgun metagenomics data from the same cohort [^3,4^] and analysed the metagenome-assembled genomes (MAGs), their distribution and functionalities when contrasted between PD and control groups. We report 952 non-redundant near-completed MAGs from 136 human gut metagenome assemblies to better understand the gut-associated microbiome genetics and diversity. We found that the *Ruminococcus bromii* genome diversity is statistically significantly higher in the control group compared to the PD group. Moreover, statistically significantly more frequently occurring genes were identified in the control group. Our results expand the knowledge of gut-associated microbiome genomes, and underscore the importance of potential consequences of strain-level differences between groups.

## Materials and methods

A summary visualisation of the methods can be seen in Supplementary Fig. S1, and Supplementary Fig. S2.

### Subjects

This study used the same patient cohort as previously described in our earlier work [^3,4^]. We included 68 control and 68 PD patients in this case-control study. See Table 1 for the characteristics of these patients.

**Table 1.** Characteristics of 136 patients included in this case-control study.

|  | Control (n=68) | PD (n=68) |
| --- | --- | --- |
| Age (years, mean ± SD) | 64.60 ± 6.87 | 65.35 ± 5.42 |
| Gender (% females) | 50% | 50% |
| BMI (kg/m<sup>2</sup>, median [IQR]) | 26.21[24.09-28.05] | 27.16[24.25-29.36] |

### Extraction of total DNA and shotgun DNA sequencing

Total DNA was extracted from frozen fecal samples using the STRATEC Molecular^®^ PSP Spin Stool DNA Plus Kit according to the manufacturer’s instructions. Nextera Library Preparation Biochemistry (Illumina) was used to prepare sequencing libraries. We performed shotgun paired-end DNA sequencing on Illumina Nextseq 500 (170 bp + 140 bp) and Illumina NovaSeq 6000 (150 bp + 150 bp) platforms.

### Read pre-processing and metagenomic assembly

Quality and adapter trimming was done using Cutadapt v1.8 [^11^] using “-m50” and truseq adapter. Reads were mapped to the Human reference genome with BWA v0.7.12-r103 [^12^], and mapped reads were filtered out. The reads were then assembled using spades v3.11.1 [^13^] with the “-meta” option.

### Binning and pangenomic analysis

We first binned the contigs using five different binning tools: Metabat v2.12.1 [^14^], MaxBin v2.2.5 [^15^], Abawaca v1.00 (https://github.com/CK7/abawaca), Concoct v0.4.0 [^16^], and MyCC [^17^] with default options. All results from the five binning tools were used as input to DASTool v1.1.0 [^18^] with the option “--search_engine diamond” to create the final MAGs. The quality of MAGs was checked using CheckM v1.0.12 [^19^]. The pangenomic analysis of selected genera was done using Anvi’o v7.0 [^20^].

### Phylogenetic analyses of MAGs

GTDB-Tk v2.1.0 (database release GTDB R07-RS207) tool [^21^] was used with the “‘classify_wf’” function to perform taxonomic annotation of each MAGs. A maximum-likelihood tree was calculated using IQ-TREE v1.6.12 [^22^] using the protein sequence alignments produced by the GTDB-Tk tool. The tree was visualised using the iTOL web tool [^23^].

### Dereplication and microdiversity profiling

We used dRep v3.2.2 [^24^] with the “dereplicate --S_algorithm fastANI --multiround_primary_clustering -ms 10000 -pa 0.9 -sa 0.95 -nc 0.30 -cm larger” options to dereplicate the constructed MAGs from all samples. To perform microdiversity analysis, each samples’ reads were mapped back to the dereplicated MAGs using Bowtie v2-2.4.4 [^25^]. Then, the inStrain v1.3.1 [^26^] tool was used with the mapping files, which also provided coverage depth of MAGs. The coverage depth data were normalised by dividing the total sample’s number of reads.

### Growth Rate Index (GRiD) calculation

We used the GRiD tool [^27^] to predict the bacterial growth rate of the dereplicated MAG set. First, a database was created using the “grid update_database” function with input of all dereplicated MAGs. Then, the “grid multiplex” function was used with the“-c 0.2” option using all the metagenomic reads per sample.

### Gene prediction, annotation and clustering

Genes from the assemblies were predicted using Prodigal [^28^] with the “-p meta” option. COG annotation of the predicted genes was done with the reCOGnizer tool [^29^]. In addition, to annotate *cas* genes we used CRISPRCasTyper [^30^]. The genes were clustered based on their nucleotide sequences with MMseqs2 [^31^] using “easy-cluster --cov-mode 1 -c 0.8 --min-seq-id 0.9” options. For amino acid-level clustering, “easy-cluster --cov-mode 1 -c 0.8 --min-seq-id 0.9” settings were used. Based on the clustering, a presence-absence table was created and Scoary v1.6.16 [^32^] was used to compare PD and control groups. For the statistically significant gene clusters, an additional detailed functional annotation was performed using PANNZER2 [^33^]. We also specifically predicted antiviral defence systems within all the MAGs using PADLOC [^34^] and PADLOC database v1.4.0 [^34^].

### Identification of viral sequences

We identified viral contigs from assemblies using VIBRANT v1.2.1 [^35^] with default settings. Viral contigs were searched within the MAGs; if a viral contig was found, we assumed the MAG is the host for the phage. The taxonomy of the identified contigs were predicted using Kaiju v1.8.2 [^36^] with Kaiju “viruses” database (downloaded on March 2022).

### Differential abundance analysis using human gut archaeome database

We used the human gut archaeome database [^37^] for archeal differential abundance analysis. Human gut archaeome database includes a few subsets based on the ANI similarity and genome completeness [^37^]. To work with a non-redundant dataset, we selected the subset with 98 archeal genomes (subset “MAGs_98” of the human gut archaeome database [^37^]). We mapped pre-processed sequencing reads to the database using Bowtie v2-2.4.4 [^25^], and the mapped reads were counted using BAMtk v0.1.1 (https://github.com/meb-team/BAM-Tk). Statistical significance between groups was calculated using DESeq2 v1.30.0 R package [^38^].

### Secondary metabolites BGCs prediction and comparison

Biosynthetic gene clusters (BGCs) putatively involved in the synthesis of secondary metabolites were predicted using antiSMASH v.5.0.0 [^39^] using full feature run as described by the developers (https://docs.antismash.secondarymetabolites.org/command_line/). Predicted BGCs were compared using automatic mode in BiG-SCAPE/CORASON [^40^] at different cut-offs (0.30, 0.40 and 0.60), and networks of similar BGCs were obtained in addition to their evolutionary histories. Selected core genes from BGCs that were not grouped with those deposited to the Minimum Information about a Biosynthetic Gene Cluster (MIBiG) repository were used to perform a BLASTp search.

## Results

### Shotgun metagenomic reads

Previously, we have observed clear differences between the PD group and the control group at the bacterial community level using the 16S rRNA gene amplicon sequencing approach [^3,4^]. In the present study, we used the same DNA samples for the shotgun metagenomics sequencing approach. Our aim was to use the metagenome data to unravel possible genes and pathways that might contribute to the PD phenotype. We focused on the metagenome-assembled genomes (MAGs), their distribution, as well as the genetic arsenal of the microbes.

In total, we analysed 136 samples; after pre-processing the sequencing reads, a total of 8,029,959,916 sequencing reads (R1+R2) were generated (Supplementary Table S1).

### Metagenomic assembly

The independent assembly of metagenomic reads of each sample showed that the total length of the assembly is larger in the control group compared to the PD group (Supplementary Fig. S3). The mean total length of the assembly was 277,969,072 bp in the control group vs. 224,912,227 bp in the PD group (only contigs larger than 1000 bp are included in the total length calculation).

### MAG reconstruction from metagenomic assembly

Assemblies were binned independently per sample using five different binning tools with DASTool aggregation (Supplementary Table S2). As a result, a total of 6,736

MAGs were retrieved, of which 3,663 resulted from the control group and 3,073 from the PD group. On average, 53 MAGs were created per sample in the control group, and 45 MAGs per sample in PD (Supplementary Fig. S3). According to CheckM results, average completeness was 86.9%, contamination was 3.7% and average genome length was 2,481,247 bp (Supplementary Table S3).

### Taxonomic annotation of MAGs

The taxonomic annotation of MAGs using the GTDB-Tk tool showed that 6,692 MAGs were annotated as bacteria, while 44 (44 of total 6,736) were archaea. On the phylum level, most (86.5%) of the bacterial MAGs were annotated as Firmicutes (synonym Bacillota) (68.1%) and Bacteroidota (synonym Bacteroidetes) (18.4%). On the genus level, the most common bacterial MAG taxa across all samples combined was *Alistipes* (299 MAGs) (Supplementary Fig. S4).

### Dereplication and MAG coverage

Since we performed independent assemblies of each sample, we encountered highly similar MAGs. To obtain non-redundant MAGs, we chose the best representative MAG using dRep [^24^]. All 6,736 MAGs were aggregated and dereplicated at 95% ANI to choose representative MAGs. In total, the dereplicated genome set consisted of 952 MAGs. Most of the MAGs were classified as bacteria (943 MAGS), while only 9 MAGs were archaea. The annotation of MAGs indicated that Firmicutes is the dominant phylum (Supplementary Fig. S5) (Figure 1). On the genus level, even with the dereplication set, we recovered 22 different versions of *Collinsella* and *Prevotella,* which indicates high variability of those genera in the human gut microbiota.

**Figure 1.**
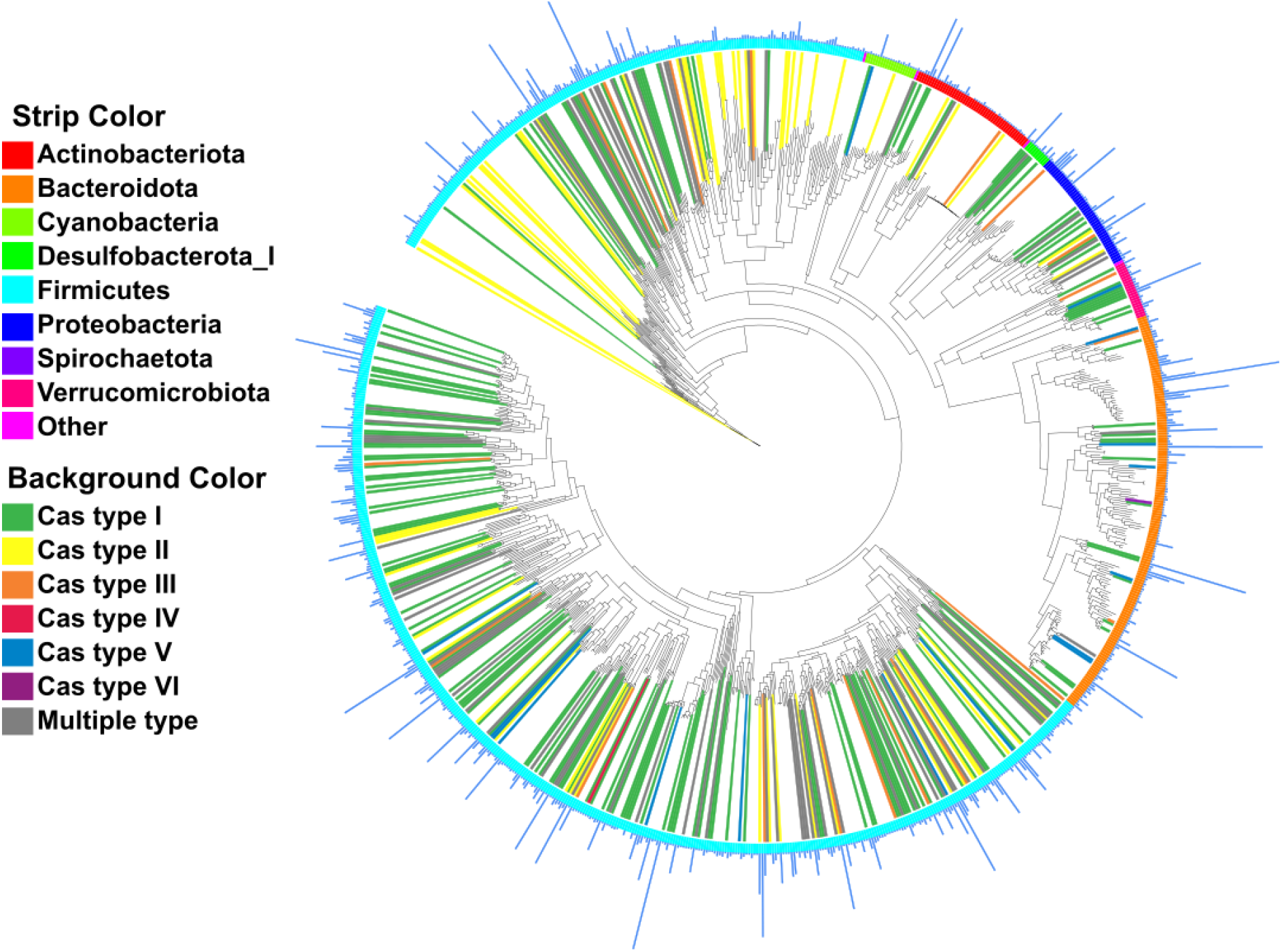
Maximum-likelihood phylogenetic tree including 943 dereplicated bacterial MAGs. Each tree branch represents a MAG. Bars on the outer layer represent how many MAGs were clustered within the dereplicated MAG, outer strip colours represent phylum of the MAGs. Background colour of the clade represents the annotated Cas type within the MAG.

The coverage depth (mean depth) of the dereplicated MAGs was calculated for each sample using the inStrain tool [^26^] by mapping back the reads to the dereplicated MAGs (Supplementary Table S4). The coverage depth data was used to understand MAG abundances across all samples. After normalising the coverage depth by sequencing depth, a Wilcoxon Rank Sum Test was applied to identify the difference in coverage depth between the control and PD groups. The results indicated that the coverage depth of 10 dereplicated MAGs was statistically significantly different between control and PD groups (please note that the significance threshold was p-adjusted to 0.1) (Supplementary Table S4, Supplementary Fig. S6). For only one MAG, coverage depth was significantly higher (p < 0.1) in the PD samples, which was taxonomically annotated as *Alistipes onderdonkii.* This suggests that the abundance of *Alistipes onderdonkii* was higher in PD samples compared to the control group. MAGs with significantly higher coverage depth (p < 0.1) in the control group belonged to genera *Agathobacter, Prevotella, Dysosmobacter, Clostridium, Choladocola,* and *Blautia* (Supplementary Fig. S6).

### Microdiversity Analysis

Microbial communities typically have different versions of bacterial genomes, presenting microdiversity that could have functional effects. Thus, reads from each sample were mapped back to the dereplicated genome set and the microdiversity profile was calculated with the inStrain tool [^26^] (Supplementary Table S4). Based on the Wilcoxon rank-sum statistics, nucleotide diversity of a MAG, annotated as *Ruminococcus bromii* B by GTDB-Tk, was statistically significantly (p = 0.002) different between the control and PD groups (Figure 2). Specifically, the nucleotide diversity was significantly higher in the control group. This indicates that *Ruminococcus bromii* is more diverse on the strain level in the control group compared to PD.

**Figure 2.**
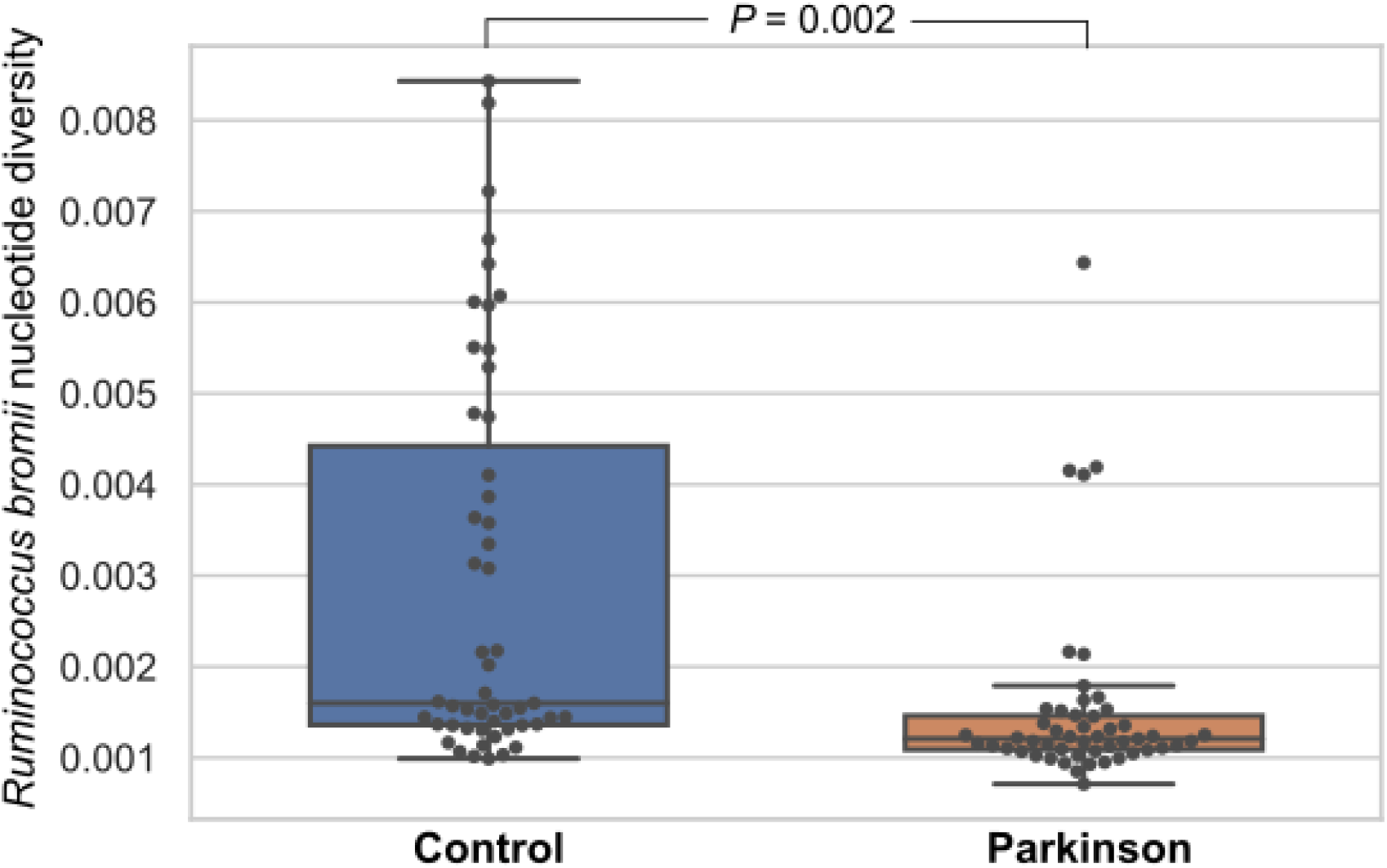
Box-plot showing nucleotide diversity of *Ruminococcus bromii* in each sample. Each dot represents one sample. Blue box represents the Control group, and the orange represents PD. The statistical significance was observed based on Wilcoxon rank-sum statistics and Benjamini/Hochberg correction.

### Growth Rate Index (GRiD)

A previous study indicated the possibility to estimate the growth of individual bacteria from shotgun metagenome data based on DNA copy number variation along the genomes [^41^]. Recently, the GRiD tool was published for making these analyses feasible for metagenomic samples [^27^]. We used this to predict the growth rate of dereplicated MAGs in each sample (Supplementary Table S5).

All samples together were used to compare growth within different phyla. Phylum-level analysis using pooled samples was done to gain understanding about the growth of a bacteria belonging to specific phylum in the gut environment in general. On average, we did not see a clear growth rate difference between microbes from different phyla. However, the data indicated some of the MAGs belonging to Firmicutes and Bacteroidota have high GRiD scores (Figure 3) compared to other phyla. The distribution of the GRiD scores also indicated that the variation is high between MAGs. For example, the minimum GRiD score within the Bacteroidota phylum was 1.00, while the maximum was 9.99 (Figure 3).

**Figure 3.**
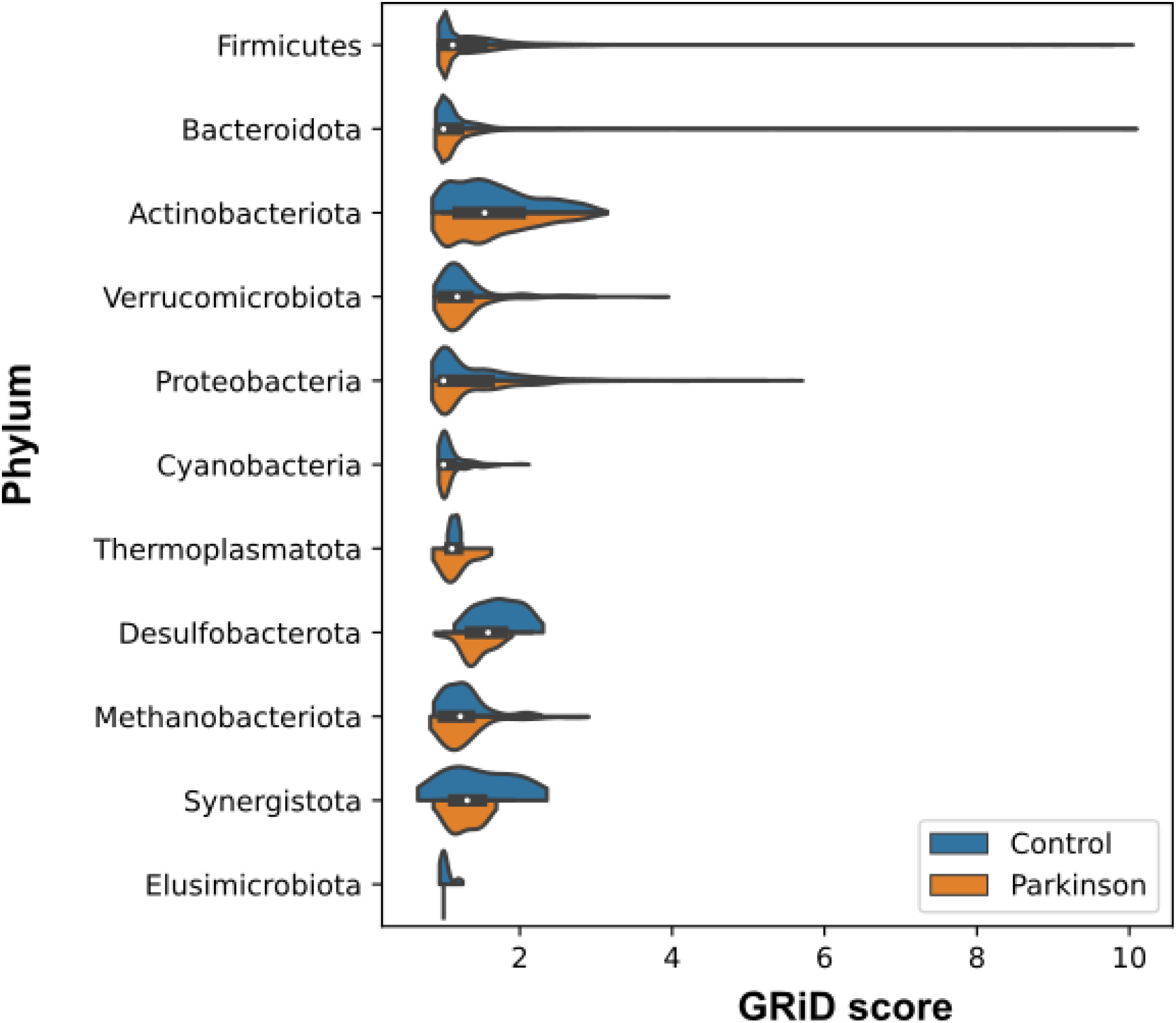
Violinplot shows the Growth Rate Index (GRiD) score of each MAGs grouped by Phylum. Blue distribution diagram represents the Control group, and orange represents PD. White dot indicates the median.

The comparison between control and PD groups showed that none of the MAGs has a statistically significant difference in growth rate. A slight but non-significant difference was observed for the growth rate (p < 0.15) for three MAGs, which were annotated as *Bifidobacterium longum, CAG-115 sp003531585* (Ruminococcaceae), and CAG-353 sp900066885 (Ruminococcaceae). For these three MAGs, the GRiD score was slightly higher in the control group (Supplementary Fig. S7).

### Pangenome of selected Genus

Previous studies showed that abundance of *Prevotella* is related to PD [^3^]. Therefore, we focused on *Prevotella* in detail for our pangenomic analysis. Pangenome analysis of 64 *Prevotella* MAGs showed that MAGs did not cluster based on the condition using average nucleotide identity (ANI) (control vs PD) (Figure 4). We also annotated genes of all *Prevotella* MAGs using the Clusters of Orthologous Genes (COG) database. We observed that the three most common COG categories in *Prevotella* MAGs were “*Cell wall/membrane/envelope biogenesis”, “Translation, ribosomal structure and biogenesis”,* and *“Carbohydrate transport and metabolism”* (Supplementary Fig. S8). To observe if any COG group was enriched in either or both PD and control groups, we used the “compute-functional-enrichment-in-pan” functionality in the Anvi’o tool [^20^]. However, no significant difference was observed between groups.

**Figure 4.**
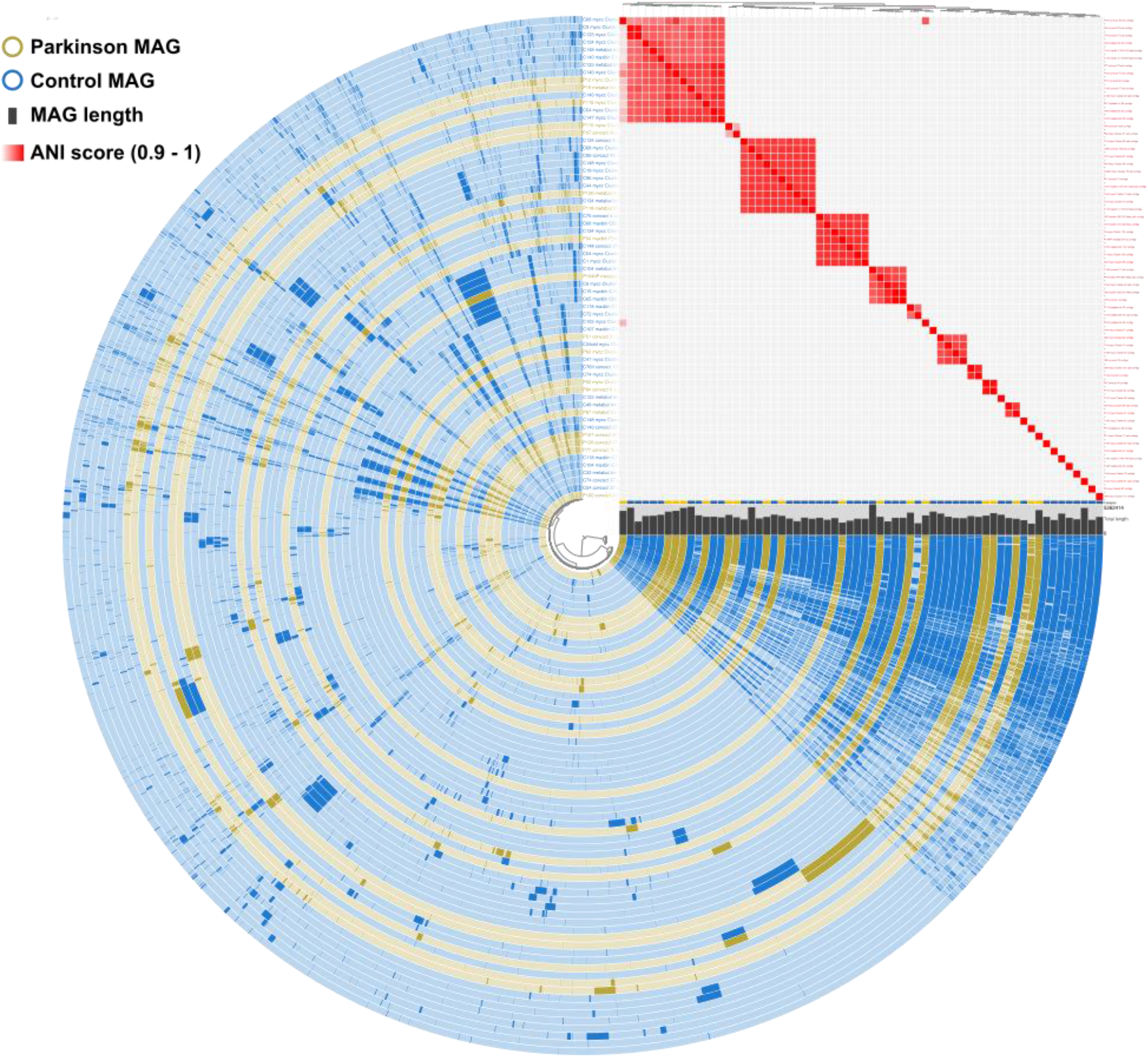
The pangenome of *Prevotella* MAGs. The dendrogram in the centre organises the 13257 gene clusters that occur in all 64 *Prevotella* MAGs. The 64 inner circular layers correspond to the 64 *Prevotella* MAG. MAGs that were created from the Control group and PD group are shown in blue and yellow, respectively. MAGs are ordered according to ANI scores which are shown at the top-right corner with red gradient. The length of the MAGs is shown with grey bars.

We additionally performed pangenomic analysis of other selected genera that were shown to be related to PD based on literature and our own studies [^3,4,42^]. However, we did not observe condition-specific differences (PD and control). COG functional enrichment was performed between the genes of control and PD MAGs for selected genera. However, we did not see any statistically significant difference in COG functions in the selected genera (Supplementary Fig. S9).

### Prediction of antiviral defence systems in MAGs

Based on PADLOC [^34^] results, 5,990 of 6,736 (89%) MAGs had at least one gene related to antiviral defence systems. For the rest of the 746 MAGs, antiviral defence system genes were not observed (Supplementary Table S6). The most common defence systems in gut microbiota were “Restriction modification type II (RM_type_II)”, “Bacterial abortive infection type E (AbiE)”, “Restriction modification type I (RM_type_I)”, and “CRISPR-Cas system type I-C (cas_type_I-C)” (Supplementary Fig. S10).

### Cas gene identification from assemblies

We also specifically identified *cas* genes from each assembly. The most common Cas operon type in gut microbiota is I-C type operon (Supplementary Fig. S11). In addition, III-A, I-E, II-A, I-B, II-C and V-A type operons were commonly detected (Supplementary Table S7) in both PD and control groups (no difference between groups for the number of Cas operon types; Supplementary Fig. S11. Similarly, we did not observe any phylum-specific cas type (Figure 1).

### Genes from metagenome assemblies

From all 136 assemblies, we predicted 93,075,923 genes, which results in an average of 684,381 predicted genes per assembly. On the MAG level, there were 15,889,713 predicted genes from the 6,736 MAGs, or an average of 2,359 genes per MAG. Since the number of predicted genes differed between assembly and MAG level, we made additional comparisons using assembly-level predictions. All 93 million genes from metagenome assemblies were annotated with the reCOGnizer (https://github.com/iquasere/reCOGnizer) tool. “Transcription” was the most common COG category, and the OmpR family DNA DNA-binding response regulator gene was the most common gene based on COG annotation (Supplementary Fig. S12, Supplementary Table S8, S9).

The predicted genes were clustered on both nucleotide and amino acid sequence-level, and the presence-absence table was constructed based on the clustering. Next, the Scoary tool [^32^] (binomial test with Benjamini–Hochberg adjustment) was used to study the association between the clustered genes’ presence or absence and the groups (control vs PD). Such analysis indicates the link between presence or absence of particular genes and observed traits (control vs PD). Statistical significance (p < 0.05) was observed for nine gene clusters on the nucleotide level, and for seven gene clusters on the amino acid level. Three of these were the same gene (Supplementary Table S10). Most of the gene clusters were seen in *Ruminococcus* and *Blautia,* and the occurrence was higher in the control group. For all significant gene clusters, the occurrence was higher in the control group compared to the PD group.

The genes with very low occurrence in PD samples can be interesting. For example, the *speF* (ornithine decarboxylase SpeF) gene from *Veillonella* (which occured in 24 control samples, but was absent in the PD samples), “Uncharacterized protein” gene from *Clostridiaceae* (occurs in 5 PD vs 34 control samples), “GIY-YIG nuclease family protein” gene from *Faecalibacterium sp.* (occurs in 1 PD sample vs 23 control samples), and “Fe-S oxidoreductase” gene from *Eubacterium sp.* (occurs in 11 PD vs 40 control samples). (Supplementary Table S10).

### Secondary metabolites biosynthetic gene clusters (BGCs)

BGCs putatively involved in the synthesis of secondary metabolites were predicted in the dereplicated MAGs using antiSMASH and compared using BiGSCAPE-CORASON (Supplementary Table S11). Some BGCs present in the MAGs were closely related to BGCs that have been previously annotated and deposited in the Minimum Information about a Biosynthetic Gene cluster (MIBiG) database (Supplementary Table S12 and Supplementary Fig. S13). The presence of biosynthetic genes involved in the synthesis of multiple molecules could be detected in MAGs dereplicated from control and PD patients, such as yersiniabactin, N-myristoyl-D-asparagine (precursor of colibactin [^43^]) and aerobactin. The BGCs present in the replicated MAGs were diverse, from which putative similar pathways were detected (Supplementary Table S11 and Supplementary Fig. S14). Core genes from the most prevalent BGCs were compared to the NCBI database (BLASTp); these genes could indicate their potential involvement in the synthesis of ranthipeptides, squalene/phytoene synthesis and arylpolyenes among biosynthetic pathways that still need to be further clarified (Supplementary Fig. S14).

### Identifying phage contigs and locating them within MAGs

From the metagenomic assemblies, we identified phages using Vibrant [^35^], which were then searched from the MAGs. If a phage (prophage) contig is located within the MAG, we assume that the MAG is the host for the phage. In total 111,099 contigs were identified as phages in the whole dataset; 31,127 of them (28%) were binned in a MAG (Supplementary Table S13). The most common phage order was Caudovirales (Supplementary Table S13). On the family level, Siphoviridae, Myoviridae, and Podoviridae were the most commonly seen phage families within the MAGs (Supplementary Table S13).

### Archeal MAGs and differential abundance analysis using human gut archaeome

From all the 6,736 identified MAGs, 44 were annotated as archaea; 26 were found in the control group, while 18 were from the PD group (Supplementary Table S14). Most of the archeal MAGs (35 of the 44) were annotated as *Methanobrevibacter smithii.* With the dereplication, we identified nine non-redundant high-quality archaeal MAGs in our complete dataset. Our coverage depth analysis showed that none of the MAGs were differentially abundant between the control and PD group.

In addition to MAGs, we also used the human gut archaeome database [^37^] for differential abundance analysis, which indicated 11 archeal genomes that differed in abundace (log2foldchange > |0.8| and p-adj < 0.05) between the control and PD group (Supplementary Table S14). Seven were statistically significantly more abundant in the PD group, six of which were annotated as *Methanobrevibacter smithii* (family level taxa is Methanobacteriaceae). Four genomes were found to be statistically significantly more abundant in the control group, all of which were annotated as Methanomethylophilaceae at the family level. Hence, our data indicated that Methanobacteriaceae was more abundant in the PD group, while Methanomethylophilaceae was more abundant in the control group.

## Discussion

Investigating the gut microbiota using shotgun sequencing and metagenome-assembled genomes (MAGs) provides more detailed genomic information compared to using only 16S RNA gene amplicon sequencing. For example, a human gut microbiota study that investigated gut microbiota MAGs during *Helicobacter pylori* eradication therapy reported that the spread of antibiotic resistance genes between

MAGs increased the relative abundance of certain MAGs [^44^]. Here we studied detailed genomic features of the MAGs that were obtained from control and PD groups. We constructed 6,736 MAGs from 136 human gut microbiota samples, corresponding to an average of 49.5 MAGs per sample. Another human gut microbiota study also indicated that on average, 53 MAGs could be created per individual sample [^44^]. With the following dereplication process, we reported 952 high-quality (min. completeness 75%, max. contamination 25%) non-redundant representative genomes from gut microbiota. Only less than 1% of the representative genomes were archaeal genomes, which is in line with a previous human gut microbiome study [^45^]. Moreover, similar to previous gut MAG studies [^45–47^], Firmicutes and Bacteroidota were the most common phyla in the gut microbiota of both the control and PD groups. On the species level, 11% of the dereplicated MAGs could not be assigned to an existing species within the GTDB database, which could indicate that those MAGs might be novel species. Similarly, Almeida et al. (2021) reported that 60% of gut metagenome MAGs could not be assigned to a species [^45^]. It is also worth mentioning that genome taxonomy databases grow parabolically; when we used Release 202 of the GTDB database (GTDB release date: April 27, 2021), 50% of MAGs were not assigned to existing species. With the Release 207 of the GTDB database (GTDB release date: April 8, 2022), only 11% of MAGs were unassigned. Here we studied quantitative differences between control and PD groups. However, it is relevant to note that part of the quantitative differences can be hampered by the MAGs building process. Reads can be disproportionately mapped to MAGs, distorting the quantitation of features like genes between the PD and control groups. Nevertheless, MAGs give a more accurate depiction of the gene and genomes compared to analysis of only reads and contigs.

Microdiversity analysis indicated that the MAGs annotated as *Ruminococcus bromii* show significant differences in strain diversity between PD and control groups. Specifically, the control group has statistically significantly (p < 0.002) more diverse *Ruminococcus bromii* genomes compared to PD. The difference in *Ruminococcus bromii* genome diversity between groups might indicate that this species adapts to the different gut microbiota environments via genomic changes. It can also be speculated that the genomic difference in *Ruminococcus* might contribute to the changes in gut microbiota environment in PD and control subjects. We observed that

*Ruminococcus* is a highly abundant genus in gut microbiota, hence genomic variation in *Ruminococcus* might influence the whole environment. Moreover, gene clustering and the presence-absence association analysis indicated that several *Ruminococcus* genes had significantly different occurrences between control and PD samples. Those genes were mostly related to metabolism, which might indicate the adaptation process.

We performed pan-genome analysis for selected genera using the generated MAGs. However, we did not see a statistically significant difference for the selected genera between control and PD groups based on COG enrichment. We then extended our gene comparison analysis to the assembly level. While about 16 million genes were predicted in the MAGs, more than 90 million genes were predicted from the assemblies of 136 individuals. Hence, by using assembly-level data, we can analyse most of the gene information in the gut environment. All predicted genes were used for clustering, and based on the cluster table, a presence or absence table was created. The statistical analysis on the presence-absence table allowed us to compare gene occurrence levels between PD and control samples. We identified some genes that were significantly more frequent in the control group, for example *speF* (encoding ornithine decarboxylase) from *Veillonella sp.* This gene cluster was not found in the PD samples, while 24 control samples harbour it. It has been reported that the ornithine level in PD patients is higher compared to controls [^48^]. It may be speculated that ornithine decarboxylase might reduce the ornithine level by converting ornithine to putrescine. The Scoary analysis indicated that several gene clusters from *Ruminococcus* and *Blautia* were significantly more frequent in the control group. As most were genes related to metabolism, one could speculate that those genes can play a role in PD protection. For example, reduced short-chain fatty acid levels have been observed in the PD group compared to the control group [^7,49,50^]. Accordingly, some fatty acid-related gene clusters were more frequent in the control group, and those genes might help to keep short-chain fatty acid levels high in the control group.

Growth rate of the gut microbes can be a useful metric to understand the growth dynamics of the gut microbial community. With the help of new tools, the replication rate of MAGs can be estimated using metagenomics data, and replication is an indication of microbial growth rate [^27^]. Our data indicated that some of the MAGs belonging to Firmicutes and Bacteroidota have a high growth rate (up to 9.9 GRiD score), which has not been seen for other phyla MAGs. This is not surprising, since Firmicutes and Bacteroidota are the most abundant phyla in gut microbiota [^45^]. The growth rate data revealed that *Bifidobacterium longum* may have a slightly higher growth rate in the control group compared to the PD group (p < 0.15). The coverage depth analysis did not indicate any difference for *Bifidobacterium longum* abundance between the control and PD group in this study. However, a higher abundance of *Bifidobacterium* has been previously reported in the PD group compared to the control group [^42^]. *Bifidobacterium* is a large genus group, and the slightly higher growth rate of *Bifidobacterium longum* in the control group here might indicate potential species-level differences in *Bifidobacterium* between the control and PD group. Hence, the whole genus pooled together shows indication of higher abundance in the PD group [^42^], while some *Bifidobacterium* species can have a higher growth rate in the control group. Therefore, some differences between groups can be only observed on the species level.

There were 10 MAGs that have a statistically significant (p < 0.1) difference in coverage depth between the control and PD group. Here, coverage depth – the average number of reads mapping to a MAG – was calculated using inStrain [^26^] by mapping the sequencing reads back to dereplicated MAGs. Coverage depth helps to understand the abundance of a microorganism, and comparing abundance between PD and control group might indicate association of a microorganism to the disease. A MAG that was annotated as *Alistipes onderdonkii* had significantly (p < 0.1) more coverage depth in the PD group, which was similarly reported in other studies with high abundance of *Alistipes* in the PD group [^51,52,53^].

In our dereplicated MAG set, there were 22 different *Collinsella* and *Prevotella* MAGs. Those had the highest number of different MAGs on the genus level. This indicates that *Collinsella* and *Prevotella* genera were highly diverse within all samples. The genus *Prevotella* is a very large group, with 717 species and 99 strains (based on NCBI Taxonomy Browser, January 2023). Previous studies showed high abundance of *Prevotella* in the control group gut microbiota compared to the PD group [^3,5,54,55^]. Similarly, we observed two *Prevotella* MAGs that had statistically significantly (p < 0.1) more coverage depth in the control group compared to the PD group, indicating higher abundance of *Prevotella* in the former (Supplementary Fig. S6). Since the MAGs were directly constructed from the patient’s gut microbiota, the abundant *Prevotella* MAGs in the control group might help future studies to focus on the *Prevotella* species. It should be noted that the coverage depth analysis (abundance analysis) was performed on the dereplicated MAG set, and further analysis using larger databases can provide more accurate abundance calculation. Such analysis was done using the same data in our other study (Pereira et al., in preparation), which also indicated higher abundance of *Prevotella* in the control group.

The dereplicated MAG set was also used to predict biosynthetic gene clusters (BGCs) that could be putatively involved in the synthesis of secondary metabolites. A previous study identified 43 BGCs that were found to be enriched in PD patients, although they were not correlated to metabolites known to be related to the disease [^56^]. Some secondary metabolite BGCs predicted from dereplicated MAGs could be involved in the synthesis of some secondary metabolites (Supplementary Table S12 and Supplementary Fig. S13). A MAG obtained from the PD sample P28 contained BGCs putatively involved in the synthesis of yersiniabactin and colibactin, which can commonly co-occur in enterobacteria [^57^]. However, the most common BGCs were present in few MAGs (up to 11 BGCs), and the produced metabolite is still unknown. Some core genes of these BGCs had sequences similar to ranthipeptides, squalene/phytoene synthesis, enterobactins, and arylpolyenes (Supplementary Fig. S14). Previous analyses based on multiomics approaches of the same cohort (but using samples taken two years later than the ones used in this study) indicated that metabolites involved in the squalene and cholesterol pathways could be used as predictive molecules [^8^], and the BGCs detected in this study could be potentially involved in the synthesis of these metabolites.

With our non-redundant MAGs, we could not detect significant differences for archeal MAGs abundance between the control and PD group. However, when we used the human gut archaeome as a database [^37^], our data indicated *Methanobrevibacter smithii* was statistically significantly (p < 0.05) more abundant in the PD group (Supplementary Table S14). This is in line with previous PD gut studies, which have previously reported that the genus *Methanobrevibacter* is more abundant [^58^] in PD patients. Recently, it has been reported that the 2-hydroxypyridine (2-HP) molecule proposed to be synthesised by the genus *Methanobrevibacter smithii* is more abundant in PD patients [^59^]. Moreover, our data indicated *Methanomethylophilaceae* UBA71 genus was statistically significantly (p < 0.05) more abundant in the control group. However, more studies are needed to understand the effect of *Methanomethylophilaceae* UBA71 in the control group.

Phages within the MAGs were analysed in this study, while further phage analyses from all metagenomic contigs are detailed in another study (Lecomte et al., in preparation). From all the 111,099 phage contigs that were identified using the 136 metagenome assemblies, 31,127 (28%) were found to be part of the predicted MAGs. Similarly, another study [^60^] reported that a host was able to be predicted for only 28.6% of all human gut phages [^60^]. We observed that the MAGs that were annotated as *Bilophila* had the highest number of phage contigs (22) within the MAGs. It should be noted that most of the predicted phage contigs are “low quality draft”, therefore the fragmented short phage contigs can be the reason for the high number of phages in *Bilophila* MAGs.

One of our interests was antiviral defence genes in the MAGs; antiviral defence systems in prokaryotes are crucial for their survival, as they are constantly infected by viruses [^61^]. Prediction of antiviral defence systems in the MAGs indicated that a very high percentage of the constructed MAGs contain at least one antiviral defence systems gene. Such a high percentage of presence of antiviral defence systems in gut bacteria might indicate that a high number of viruses can be found in gut microbiota, and gut bacteria adapt well to defend themselves from viruses. To our knowledge, the complete antiviral defence system search has not been performed for other metagenomic environmental samples, therefore it is not known if the high presence of antiviral defence systems is specific to gut microbiota or more widespread.

## Supporting information

All supplementary figures and tables

## Acknowledgements

The authors would like to thank the personnel of the DNA Sequencing and Genomics Laboratory for running the NGS assays. We acknowledge the CSC - IT Center for Science, Finland, for computational resources, and the University of Helsinki Language Services for English language revision.

## Author contributions

F.S., P.A.B.P., and P.A. conceived and designed the study. F.S. performed clinical evaluation of the patients. P.A.B.P gave statistical support. I.C.D, A.L., T.K.S., P.L., and J.S. analysed the sequencing data. L.P organised NGS assays. P.A. and I.C.D. drafted the manuscript. All authors reviewed the manuscript.

## Data availability

All sequencing data and constructed MAGs have been deposited in the European Nucleotide Archive (ENA) under accession code PRJEB59350.

