## Supplementary material for "Metagenome-assembled microbial genomes from Parkinson’s disease fecal samples": All supplementary figures and tables: Supplemental_TableS12.docx

Table S12. Gene cluster families that were phylogenetically grouped with biosynthetic gene clusters (BGC) deposited in the Minimum Information about a Biosynthetic Gene cluster (MIBiG). Information obtained using BiGSCAPE/CORASON in cutoff 0.60.

| **Type** | **MIBiG (organism studied)** | **Compound (from MIBiG)** | **Parkinson disease** | **Control** | **Gene cluster family** |
| --- | --- | --- | --- | --- | --- |
| NRPS | BGC0001575.1 (*Ruminococcus* sp.) | dipeptide aldehydes | c00078_P66  c00026_P47  c00007_P105 | c00030_C28 | FAM_01479 |
| NRPS | BGC0001686.1 (*Clostridium* sp.) | N-octanoyl-Met-Phe-H | c00009_P47  c00011_P119  c00008_P45 | c00028_C72 | FAM_02167 |
| NRPS, PKSother, PKS-NRPS | BGC0001055.1 (*Escherichia coli*)  BGC0000467.1 (*Yersinia Enterocolitica*) | yersiniabactin | c00032_P28  c00032_P63 |  | FAM_00445 (FAM_02411) |
| NRPS, PKSother, PKS-NRPS | BGC0000972.1 (*Escherichia coli*) | N-myristoyl-D-asparagine | c00028_P28 |  | FAM_02386 |
| RiPPs | BGC0000624.1 (*Lactobacillus salivarius*) | salivaricin CRL1328 α/β peptide | c00033_P70 |  | FAM_00597 |
| RiPPs | BGC0001229.1 (*Streptococcus bingchenggensis*)  BGC0001311.1 (*Ruminococcus flavefaciens*)  BGC0001210.1 (*Bacillus pseudomycoides*)  BGC0000554.1 (*Streptomyces filamentosus*) | bingicin α/β, flavecins, pseudomycoicidin, SRO15-3108 (lanthipeptide) |  | c00084_C68 | FAM_01150 |
| RiPPs | BGC0001209.1 (*Streptococcus thermophilus*)  BGC0001929.1 (*Streptococcus ferus*) | Streptide, WGK | c00066_P19 |  | FAM_01149 |
| RiPPs | BGC0001602.1 (*Lactococcus gasseri*)  BGC0000619.1 (*Lactococcus gasseri*)  BGC0001388.1 (*Lactococcus gasseri*) | gassericin-T/E | c00027_P103 | c00017_C76II  c00001_C70 | FAM_01298 |
| RiPPs | BGC0000534.1 (*Streptococcus mutans*)  BGC0000557.1 (*Streptococcus pyogenes*)  BGC0001788.1 (*Streptococcus suis*)  BGC0000526.1 (*Streptococcus macedonicus*)  BGC0000547.1 (*Streptococcus salivari*)  BGC0000521.1 (*Lactococcus lactis*)  BGC0000539.1 (*Staphylococcus warneri*)  BGC0000545.1 (*Ruminococcus gnavus*) | Mutation K8, streptococcin A-FF22, suicin 65, macedocin, salivaricin 9, lacticin481, nukacin ISK-1, ruminococcin A (lantibiotic) |  | c00003_C20 | FAM_02014 |
| Others | BGC0000836.1 (*Escherichia coli*) | APE Ec (*E. coli* aryl polyene) | c00062_P28 | c00151_C34old | FAM_02522 |
| Others | BGC0002008.1 (*Xenorhabdus doucetiae*)  BGC0000837.1 (*Vibrio fischeri*) | Aryl polyenes | c00024_P69  c00004_P115 | c00022_C9  c00014_C70  c00002_C44 | FAM_01887 |
| Others | BGC0000839.1 (*Chitinophaga pinensis*) | Flexirubin | c00092_P69  c00064_P115  c00014_P51  c00005_P4  c00018_P68  c00037_P71 | c00123_C147  c00001_C80  c00020_C18 | FAM_02526 |
| Others | BGC0001498.1 (*Xenorhabdus szentirmaii*)  BGC0001499.1 (*Pantoea ananatis*) BGC0001555.1 (*Escherichia coli*)  BGC0000939.1 (*Grimontia hollisae*) | aerobactin, colicin V |  | c00026_C44  c00046_C9 | FAM_01403 |
