## Supplementary material for "Metagenome-assembled microbial genomes from Parkinson’s disease fecal samples": All supplementary figures and tables: Supplementary_FigureS6.pdf

MAG: C82.metabat.bin.11.contigs
Taxa: Alistipes onderdonkii

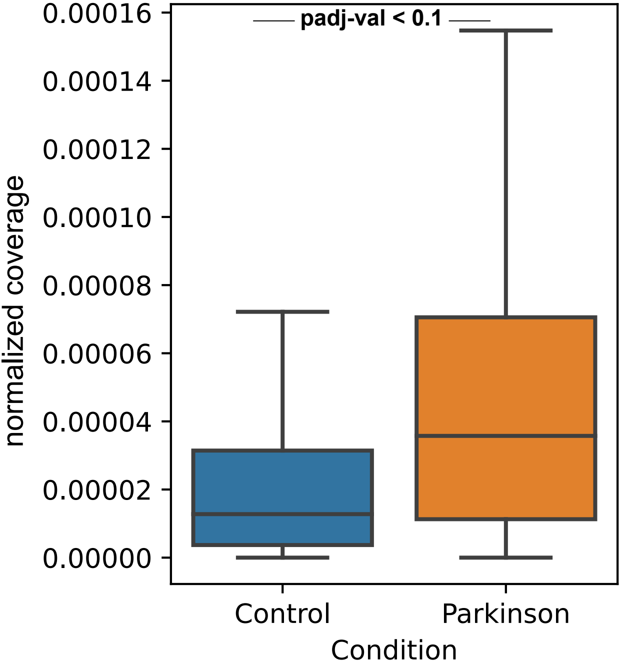

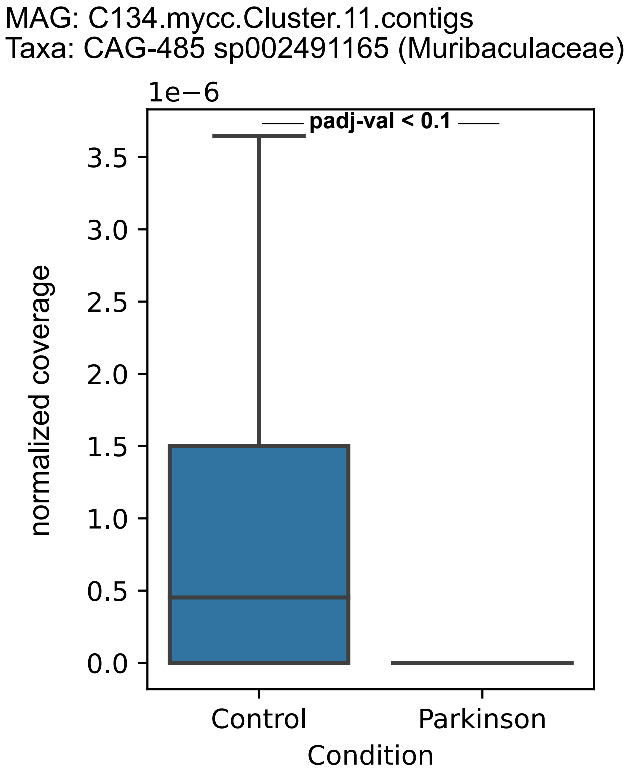

MAG: C134.mycc.Cluster.11.contigs
Taxa: CAG-485 sp002491165 (Muribaculaceae)
1e−6
padj-val < 0.1
normalized coverage
0.0
0.5
1.0
1.5
2.0
2.5
3.0
3.5
Control
Parkinson
Condition

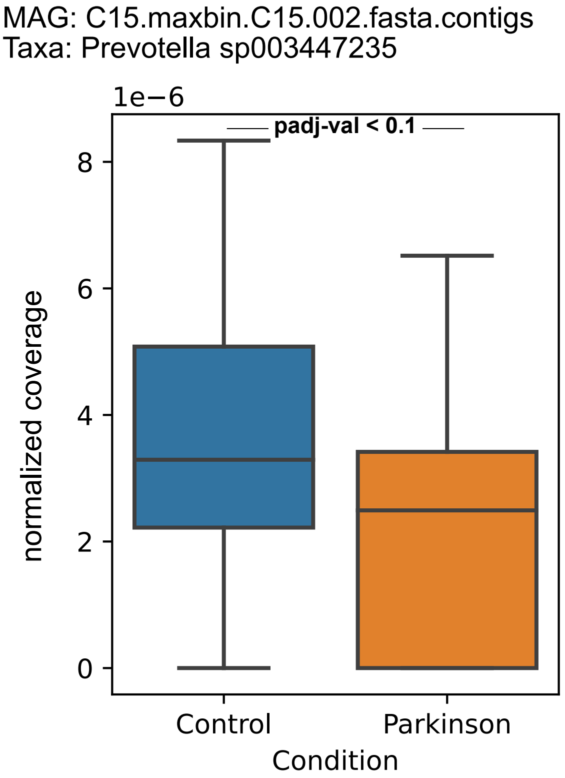

MAG: C15.maxbin.C15.002.fasta.contigs
Taxa: Prevotella sp003447235
1e−6
padj-val < 0.1
8
6
4
2
0
normalized coverage
Control
Parkinson
Condition

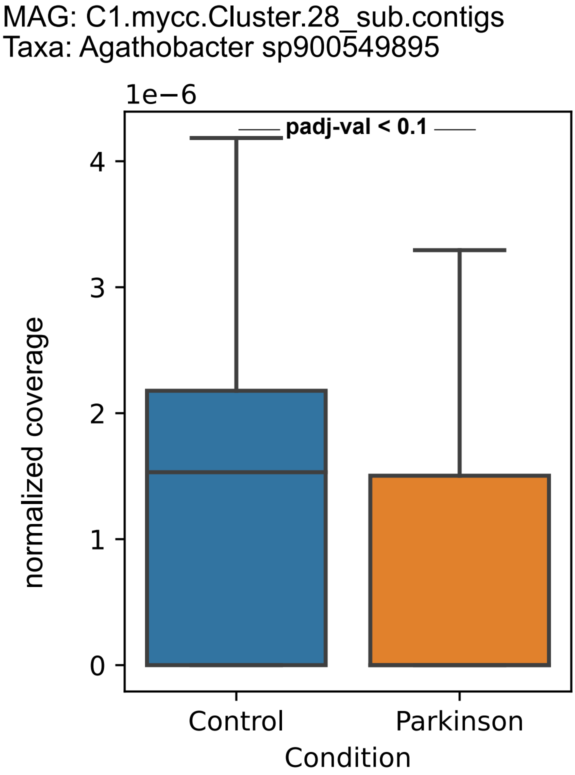

MAG: C1.mycc.Cluster.28_sub.contigs
Taxa: Agathobacter sp900549895
1e−6
padj-val < 0.1
0
1
2
3
4
normalized coverage
Control
Parkinson
Condition

MAG: C33.metabat.bin.54_sub.contigs
Taxa: Agathobacter sp000434275

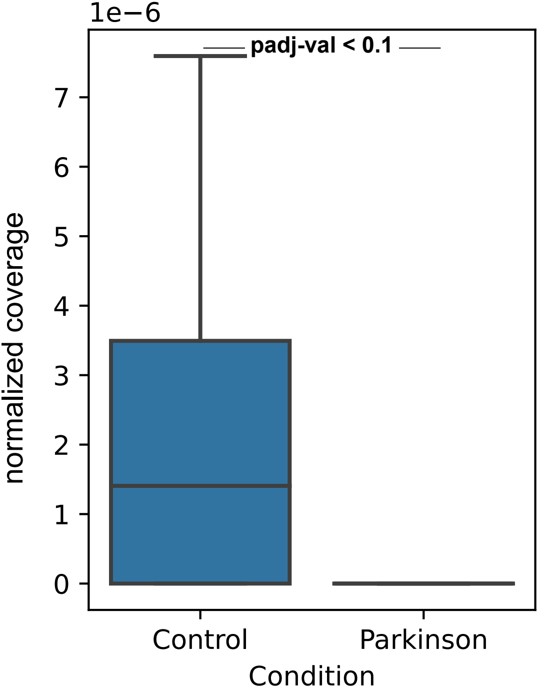

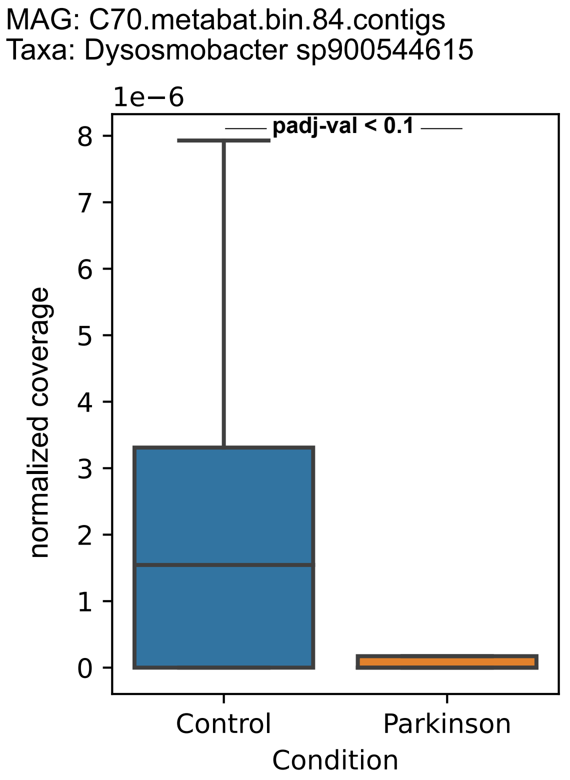

MAG: C70.metabat.bin.84.contigs
Taxa: Dysosmobacter sp900544615
1e−6
padj-val < 0.1
8
7
6
5
4
3
2
1
0
normalized coverage
Control
Parkinson
Condition

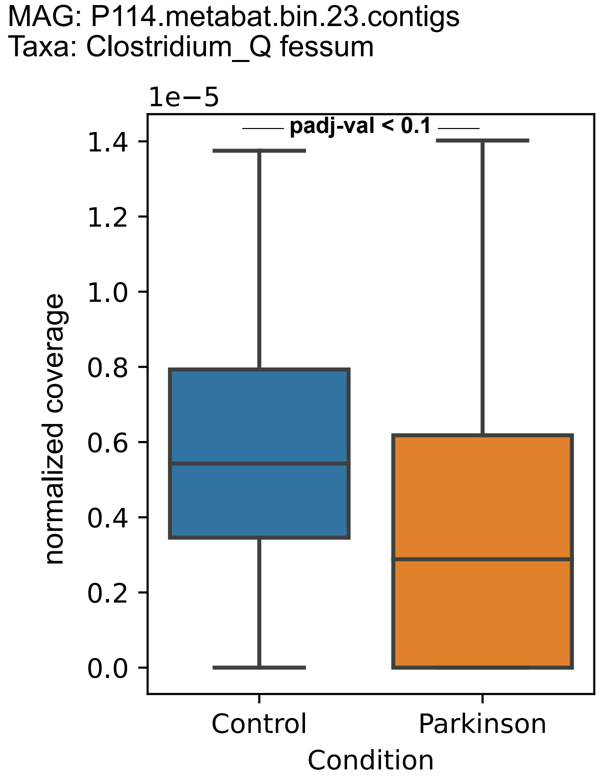

MAG: P114.metabat.bin.23.contigs
Taxa: Clostridium_Q fessum
1e−5
padj-val < 0.1
1.4
1.2
1.0
0.8
0.6
0.4
0.2
0.0
normalized coverage
Control
Parkinson
Condition

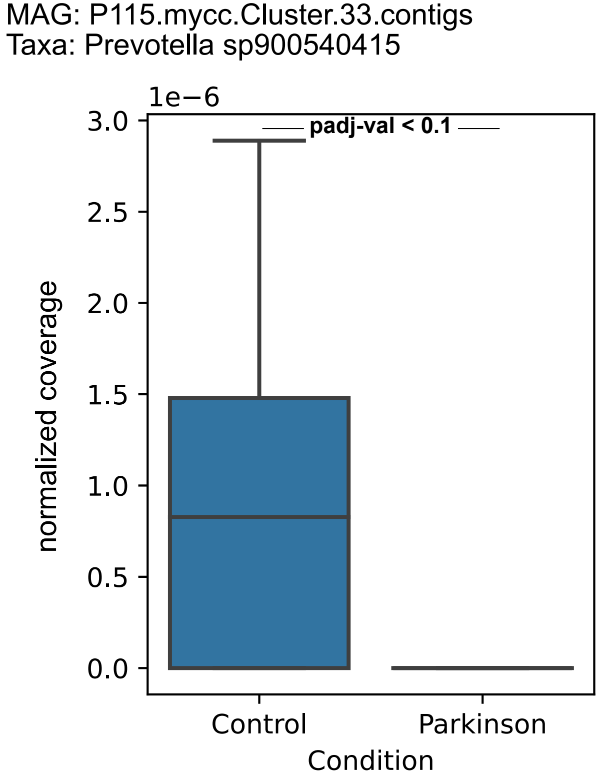

MAG: P115.mycc.Cluster.33.contigs
Taxa: Prevotella sp900540415
1e−6
padj-val < 0.1
3.0
2.5
2.0
1.5
1.0
0.5
0.0
normalized coverage
Control
Parkinson
Condition

MAG: P16.metabat.bin.41.contigs
Taxa: Choladocola sp018223365

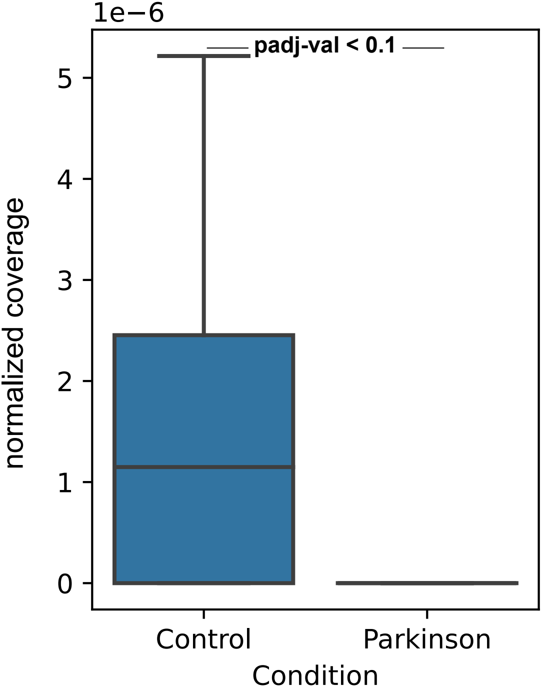

MAG: P66.metabat.bin.18_sub.contigs
Taxa: Blautia_A wexlerae

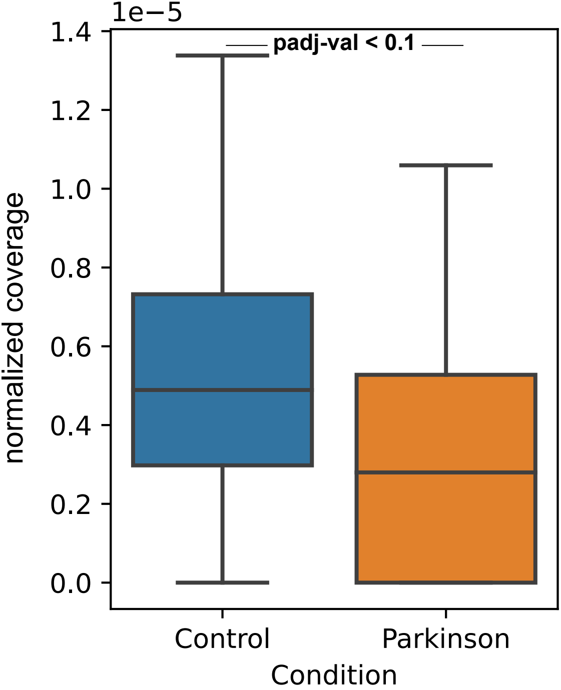

Figure S6. Boxplot of each MAGs that we observed statistically significant difference between two groups in coverage data. The coverage is normalized by the whole sequencing depth. Blue box represent Control samples and orange Parkinson samples. Statistical difference between groups were calculated using Wilcoxon rank-sum statistic with Benjamini/Hochberg correction.
