## Supplementary material for "Metagenome-assembled microbial genomes from Parkinson’s disease fecal samples": All supplementary figures and tables: Supplementary_FigureS7.pdf

**MAG: P115.metabat.bin.75.contigs**
**GTDB-tk annotation: *Bifidobacterium longum***

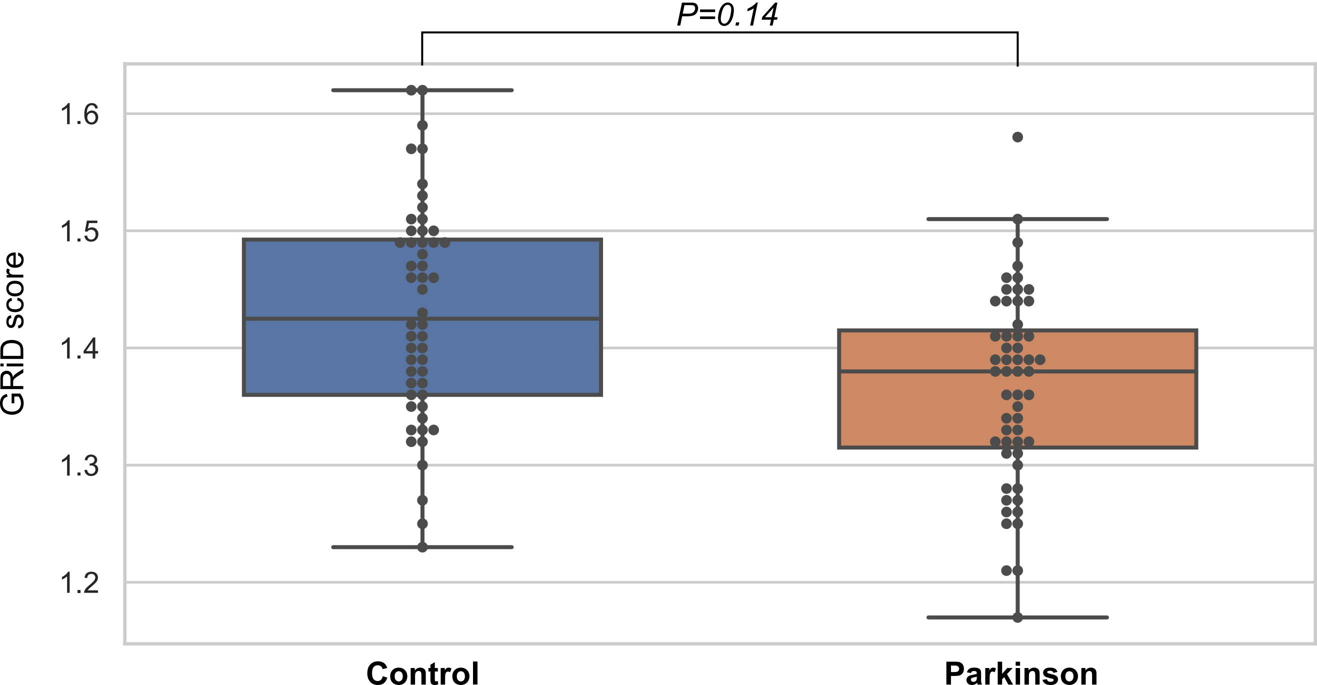

**MAG: P60.maxbin.P60.030.fasta_sub.contigs**
**GTDB-tk annotation: CAG-115 sp003531585 (*Ruminococcaceae*)**

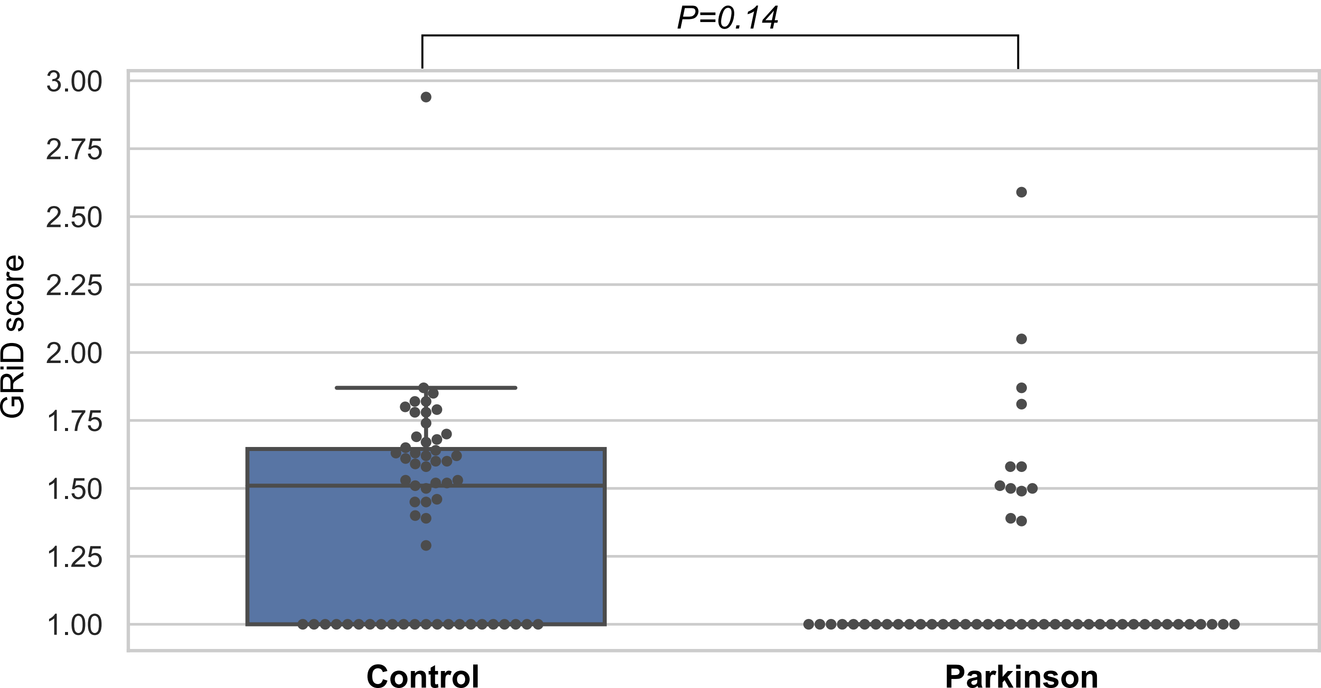

MAG: C18.mycc.Cluster.92.contigs
GTDB-tk annotation: CAG-353 sp900066885 (*Ruminococcaceae*)

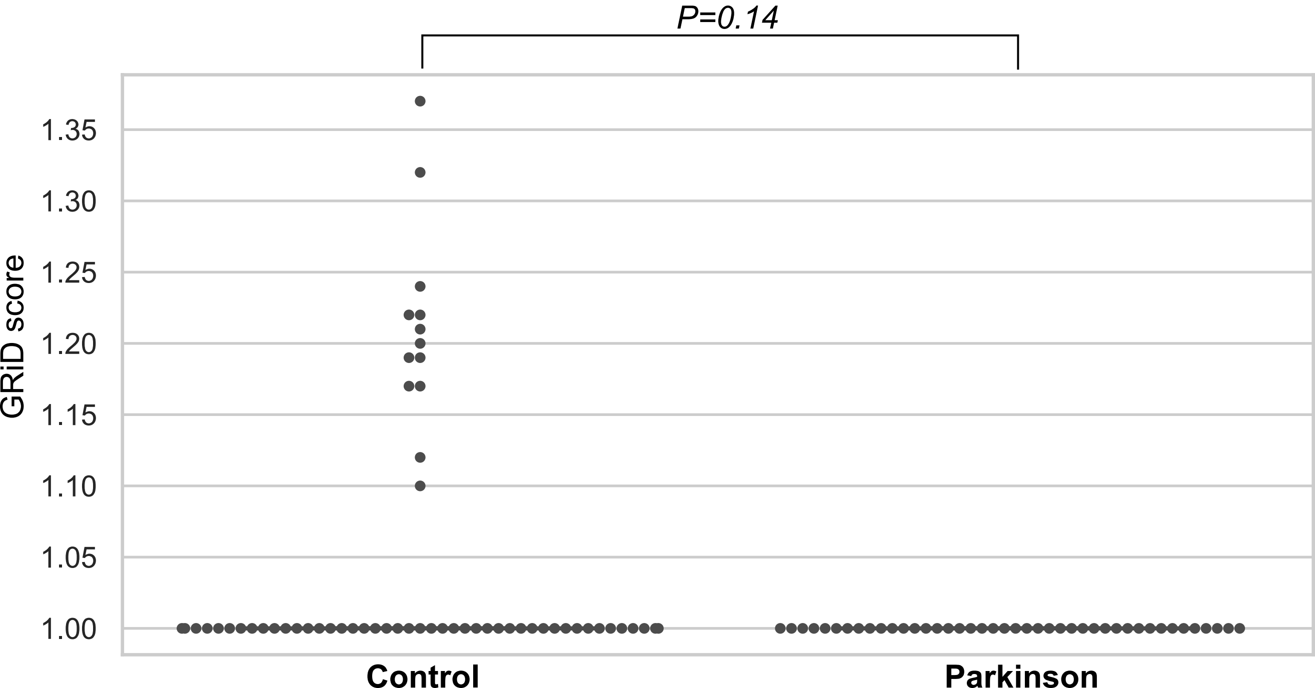

Supplementary Figure S7. Boxplot of each MAGs that we observed difference between two groups with low P-value in Growth Rate Index (GRiD) score. Each dot represents one sample. Blue box represents the Control group, and orange Parkinson. Statistical difference between groups were calculated using Wilcoxon rank-sum statistic with Benjamini/Hochberg correction.
