## Supplementary material for "Metagenome-assembled microbial genomes from Parkinson’s disease fecal samples": All supplementary figures and tables: Supplementary_FigureS8.pdf

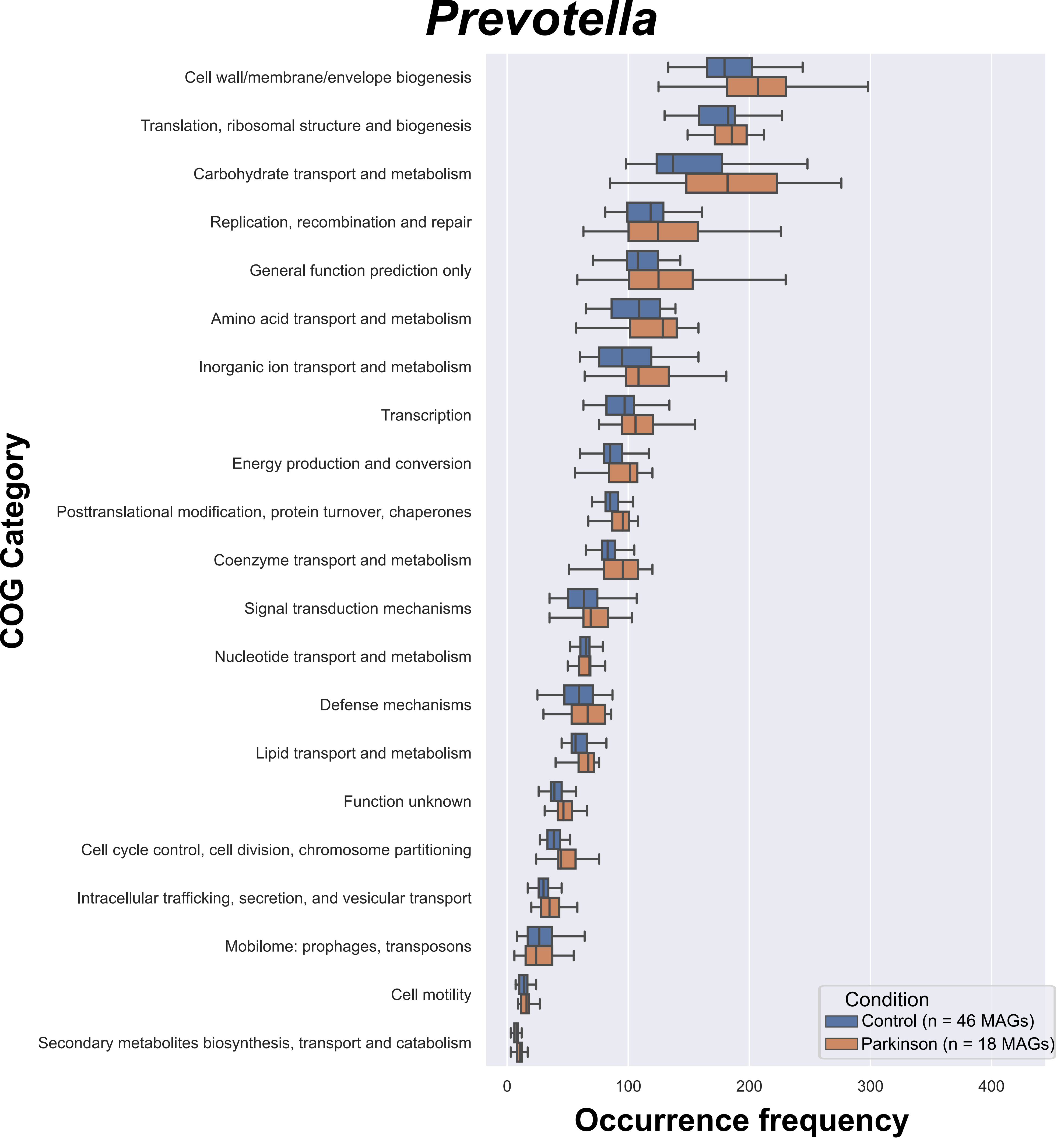


Figure S8. The occurrence frequencies of COG categories in Prevotella MAGs. Each COG category is represented on y axis, and box plot represents the occurrence frequency within the MAGs. Blue is control MAGs, and orange is Parkinson MAGs.
