## Supplementary material for "Metagenome-assembled microbial genomes from Parkinson’s disease fecal samples": All supplementary figures and tables: Supplementary_FigureS9.pdf

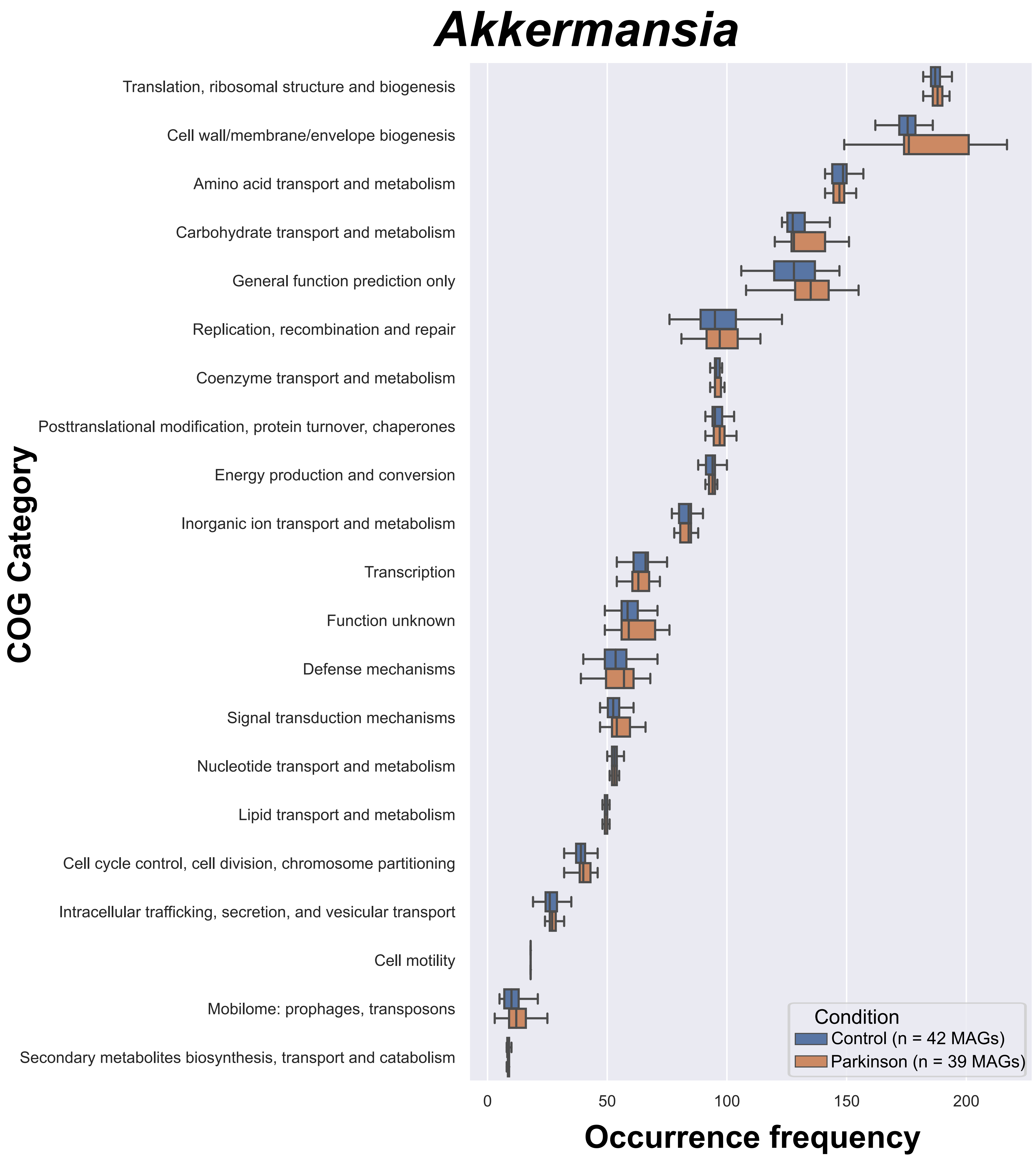

Akkermansia
COG Category
Translation, ribosomal structure and biogenesis
Cell wall/membrane/envelope biogenesis
Amino acid transport and metabolism
Carbohydrate transport and metabolism
General function prediction only
Replication, recombination and repair
Coenzyme transport and metabolism
Posttranslational modification, protein turnover, chaperones
Energy production and conversion
Inorganic ion transport and metabolism
Transcription
Function unknown
Defense mechanisms
Signal transduction mechanisms
Nucleotide transport and metabolism
Lipid transport and metabolism
Cell cycle control, cell division, chromosome partitioning
Intracellular trafficking, secretion, and vesicular transport
Cell motility
Mobilome: prophages, transposons
Secondary metabolites biosynthesis, transport and catabolism
Condition
Control (n = 42 MAGs)
Parkinson (n = 39 MAGs)
0
50
100
150
200
Occurrence frequency

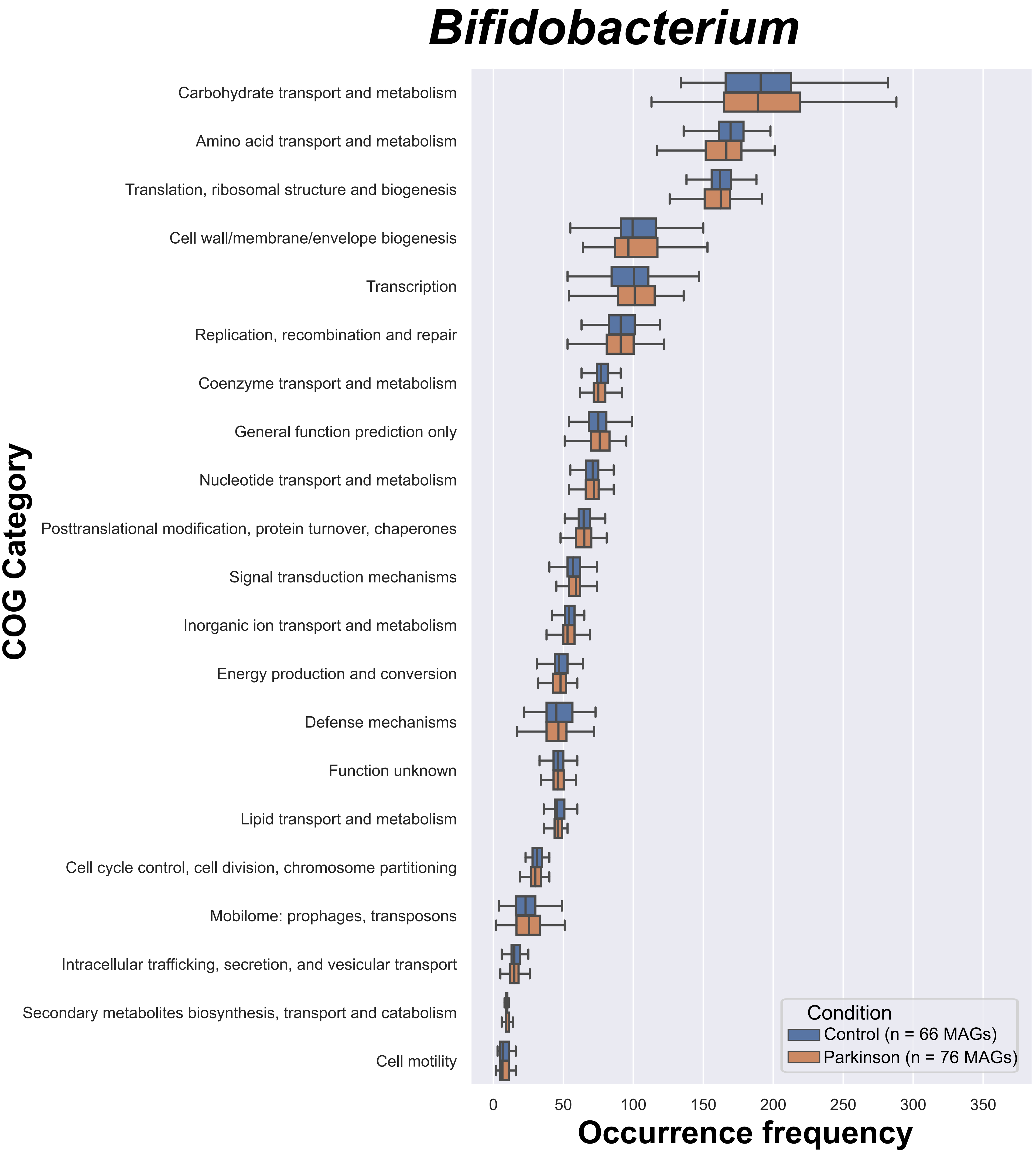
Bifidobacterium
COG Category
Carbohydrate transport and metabolism
Amino acid transport and metabolism
Translation, ribosomal structure and biogenesis
Cell wall/membrane/envelope biogenesis
Transcription
Replication, recombination and repair
Coenzyme transport and metabolism
General function prediction only
Nucleotide transport and metabolism
Posttranslational modification, protein turnover, chaperones
Signal transduction mechanisms
Inorganic ion transport and metabolism
Energy production and conversion
Defense mechanisms
Function unknown
Lipid transport and metabolism
Cell cycle control, cell division, chromosome partitioning
Mobilome: prophages, transposons
Intracellular trafficking, secretion, and vesicular transport
Secondary metabolites biosynthesis, transport and catabolism
Cell motility
0
50
100
150
200
250
300
350
Occurrence frequency
Condition
Control (n = 66 MAGs)
Parkinson (n = 76 MAGs)

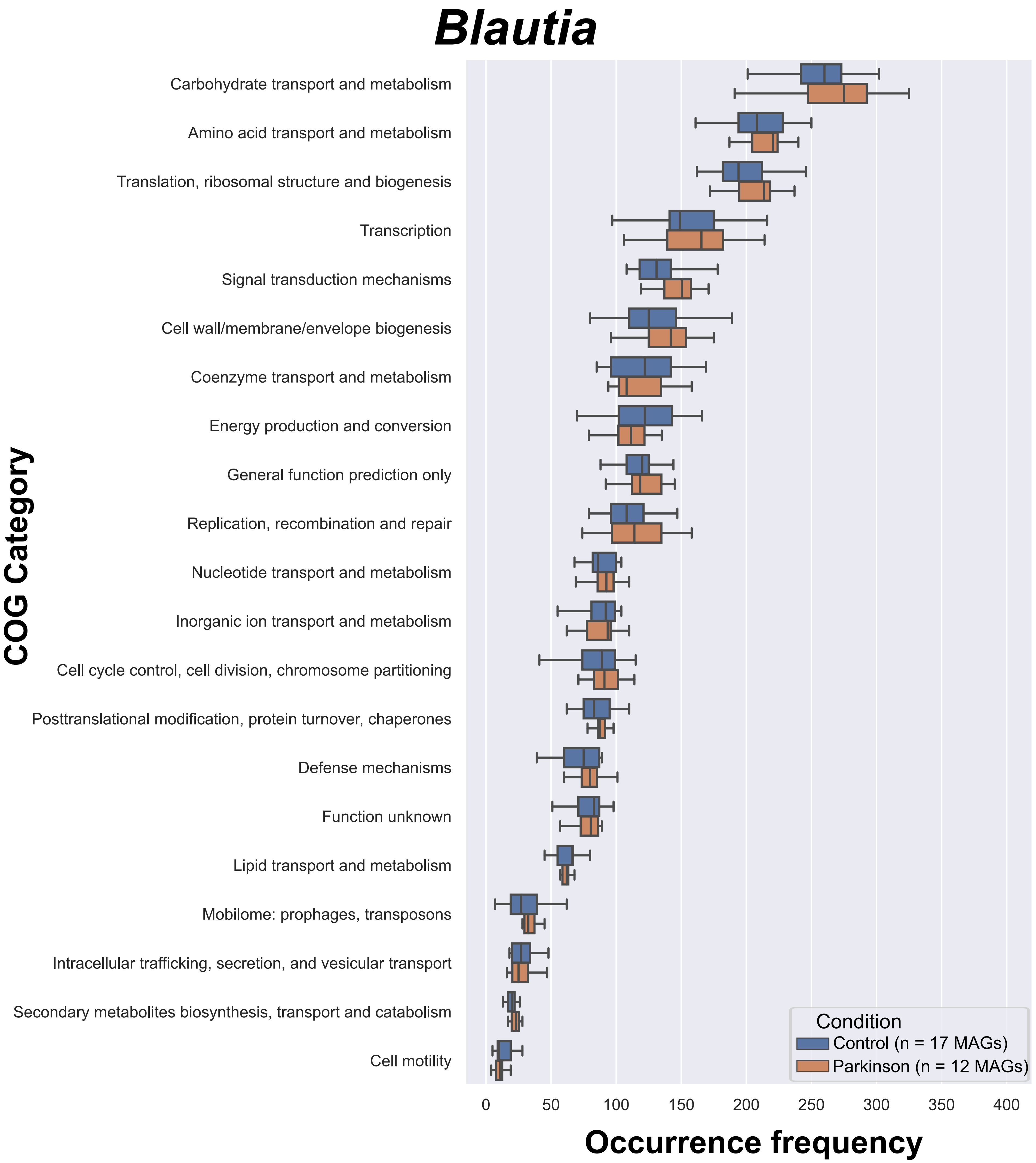
Blautia
COG Category
Carbohydrate transport and metabolism
Amino acid transport and metabolism
Translation, ribosomal structure and biogenesis
Transcription
Signal transduction mechanisms
Cell wall/membrane/envelope biogenesis
Coenzyme transport and metabolism
Energy production and conversion
General function prediction only
Replication, recombination and repair
Nucleotide transport and metabolism
Inorganic ion transport and metabolism
Cell cycle control, cell division, chromosome partitioning
Posttranslational modification, protein turnover, chaperones
Defense mechanisms
Function unknown
Lipid transport and metabolism
Mobilome: prophages, transposons
Intracellular trafficking, secretion, and vesicular transport
Secondary metabolites biosynthesis, transport and catabolism
Cell motility
Condition
Control (n = 17 MAGs)
Parkinson (n = 12 MAGs)
0
50
100
150
200
250
300
350
400
Occurrence frequency

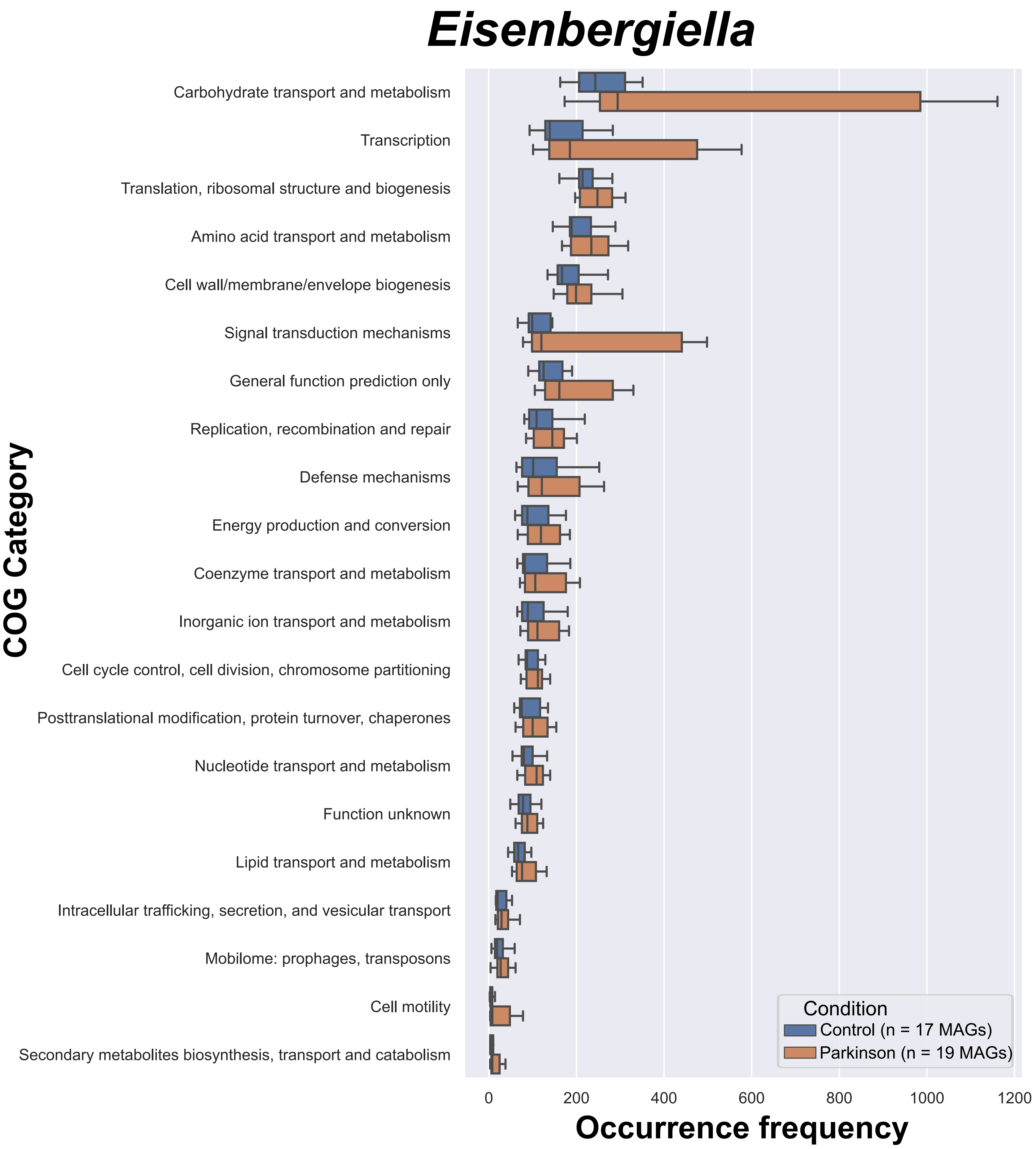
Eisenbergiella
COG Category
Carbohydrate transport and metabolism
Transcription
Translation, ribosomal structure and biogenesis
Amino acid transport and metabolism
Cell wall/membrane/envelope biogenesis
Signal transduction mechanisms
General function prediction only
Replication, recombination and repair
Defense mechanisms
Energy production and conversion
Coenzyme transport and metabolism
Inorganic ion transport and metabolism
Cell cycle control, cell division, chromosome partitioning
Posttranslational modification, protein turnover, chaperones
Nucleotide transport and metabolism
Function unknown
Lipid transport and metabolism
Intracellular trafficking, secretion, and vesicular transport
Mobilome: prophages, transposons
Cell motility
Secondary metabolites biosynthesis, transport and catabolism
Condition
Control (n = 17 MAGs)
Parkinson (n = 19 MAGs)
0
200
400
600
800
1000
1200
Occurrence frequency

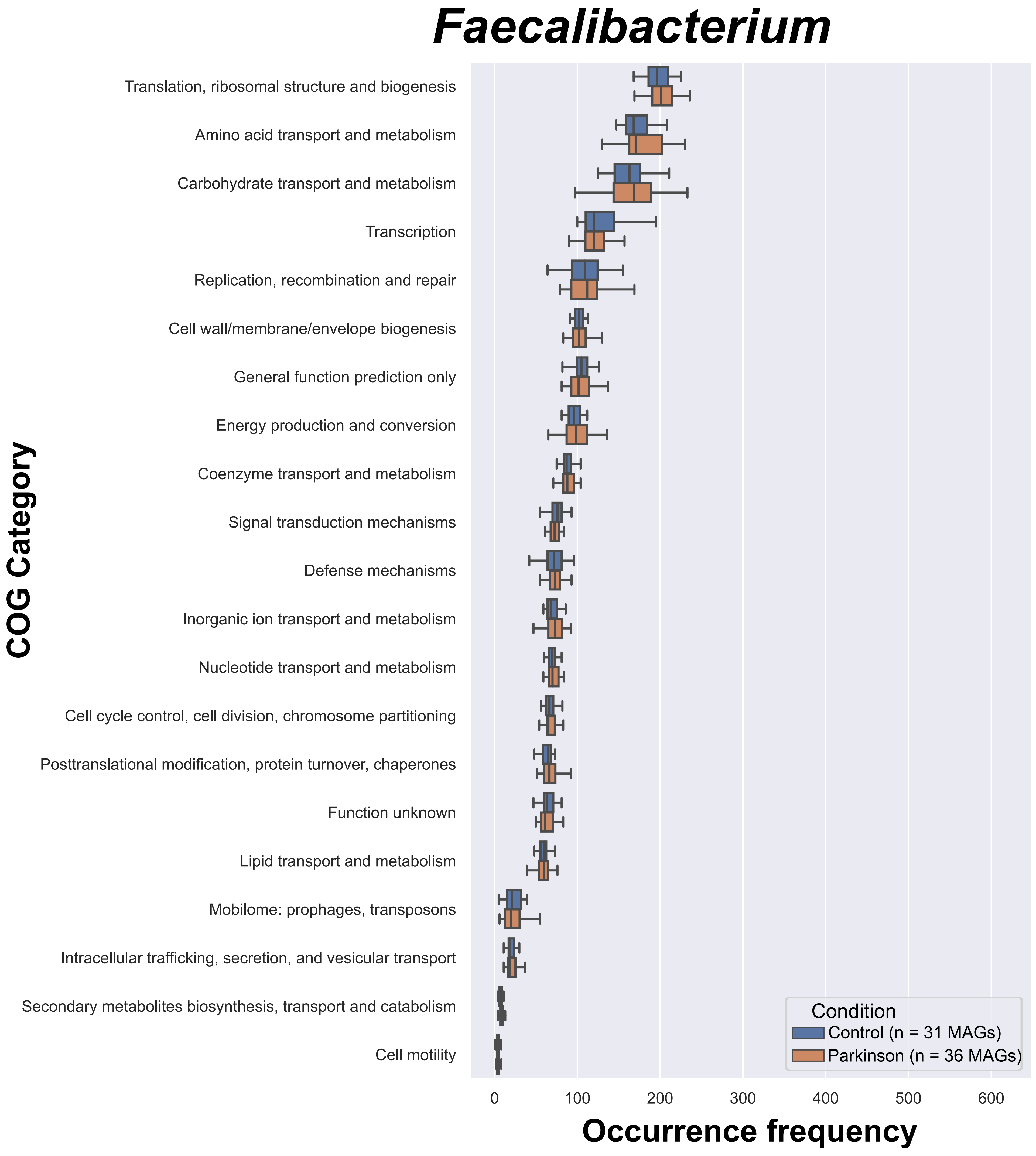
Faecalibacterium
COG Category
Translation, ribosomal structure and biogenesis
Amino acid transport and metabolism
Carbohydrate transport and metabolism
Transcription
Replication, recombination and repair
Cell wall/membrane/envelope biogenesis
General function prediction only
Energy production and conversion
Coenzyme transport and metabolism
Signal transduction mechanisms
Defense mechanisms
Inorganic ion transport and metabolism
Nucleotide transport and metabolism
Cell cycle control, cell division, chromosome partitioning
Posttranslational modification, protein turnover, chaperones
Function unknown
Lipid transport and metabolism
Mobilome: prophages, transposons
Intracellular trafficking, secretion, and vesicular transport
Secondary metabolites biosynthesis, transport and catabolism
Cell motility
Condition
Control (n = 31 MAGs)
Parkinson (n = 36 MAGs)
0
100
200
300
400
500
600
Occurrence frequency

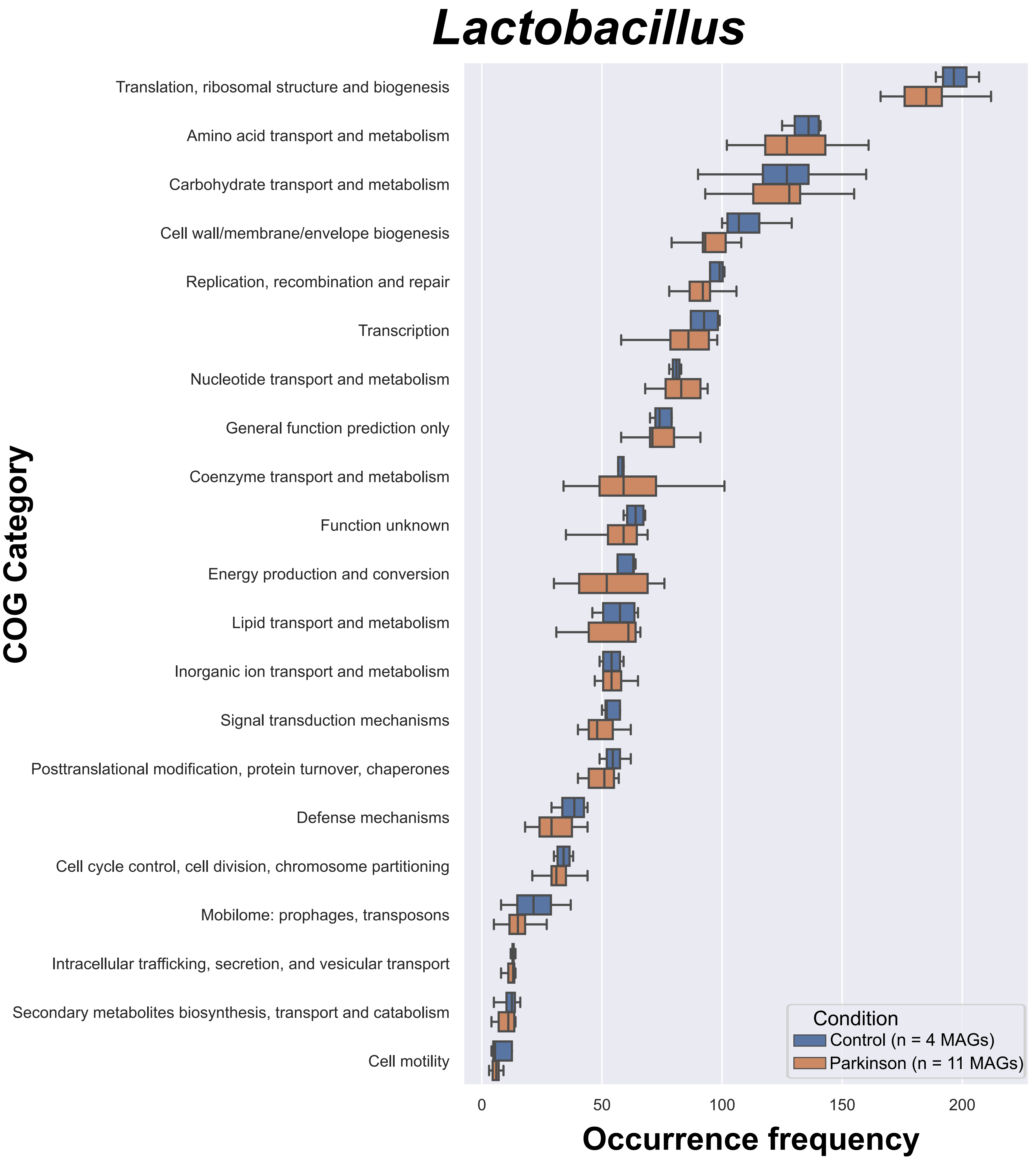
Lactobacillus
COG Category
Translation, ribosomal structure and biogenesis
Amino acid transport and metabolism
Carbohydrate transport and metabolism
Cell wall/membrane/envelope biogenesis
Replication, recombination and repair
Transcription
Nucleotide transport and metabolism
General function prediction only
Coenzyme transport and metabolism
Function unknown
Energy production and conversion
Lipid transport and metabolism
Inorganic ion transport and metabolism
Signal transduction mechanisms
Posttranslational modification, protein turnover, chaperones
Defense mechanisms
Cell cycle control, cell division, chromosome partitioning
Mobilome: prophages, transposons
Intracellular trafficking, secretion, and vesicular transport
Secondary metabolites biosynthesis, transport and catabolism
Cell motility
Condition
Control (n = 4 MAGs)
Parkinson (n = 11 MAGs)
0
50
100
150
200
Occurrence frequency

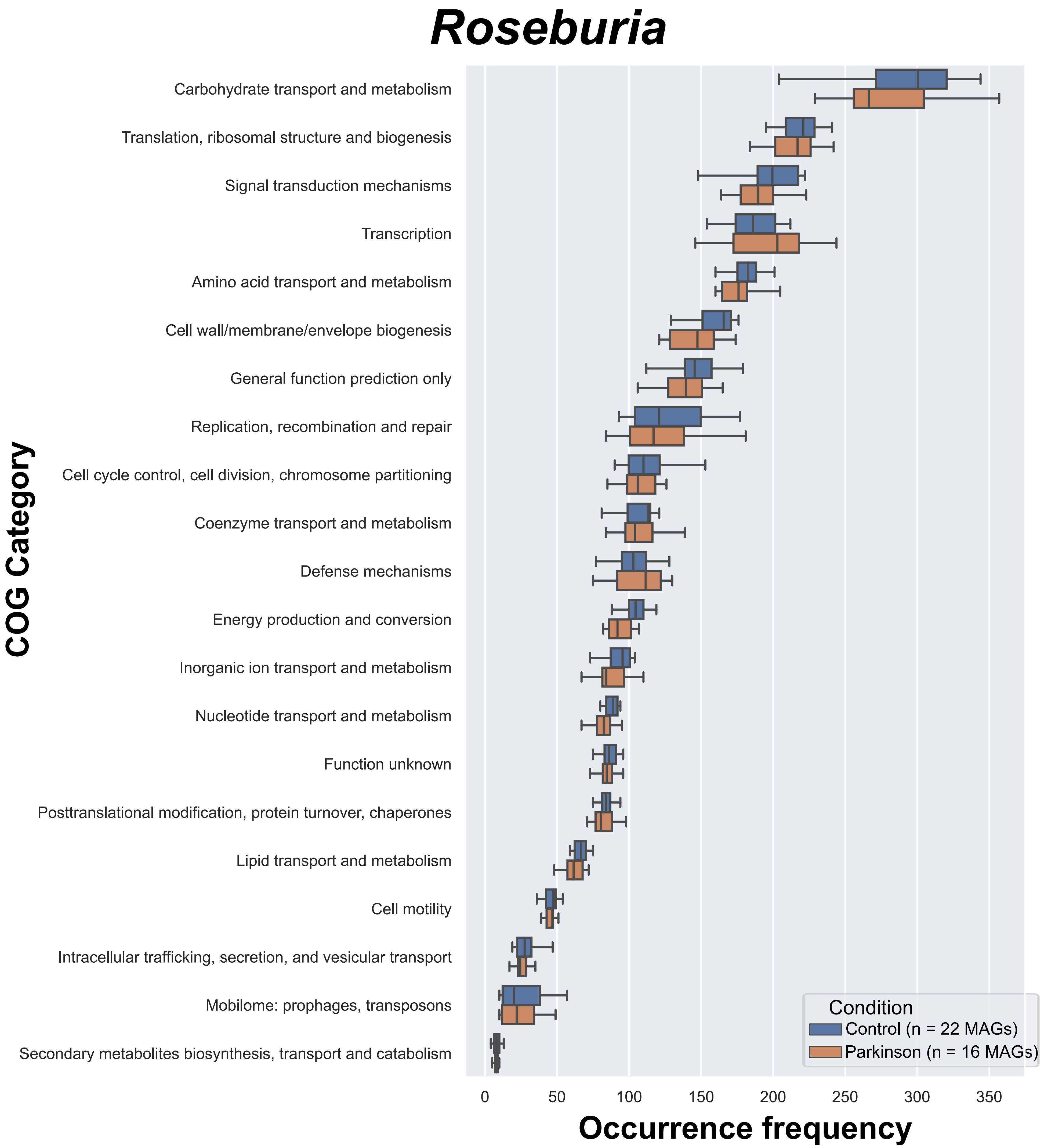
Roseburia
COG Category
Carbohydrate transport and metabolism
Translation, ribosomal structure and biogenesis
Signal transduction mechanisms
Transcription
Amino acid transport and metabolism
Cell wall/membrane/envelope biogenesis
General function prediction only
Replication, recombination and repair
Cell cycle control, cell division, chromosome partitioning
Coenzyme transport and metabolism
Defense mechanisms
Energy production and conversion
Inorganic ion transport and metabolism
Nucleotide transport and metabolism
Function unknown
Posttranslational modification, protein turnover, chaperones
Lipid transport and metabolism
Cell motility
Intracellular trafficking, secretion, and vesicular transport
Mobilome: prophages, transposons
Secondary metabolites biosynthesis, transport and catabolism
0
50
100
150
200
250
300
350
Occurrence frequency
Condition
Control (n = 22 MAGs)
Parkinson (n = 16 MAGs)

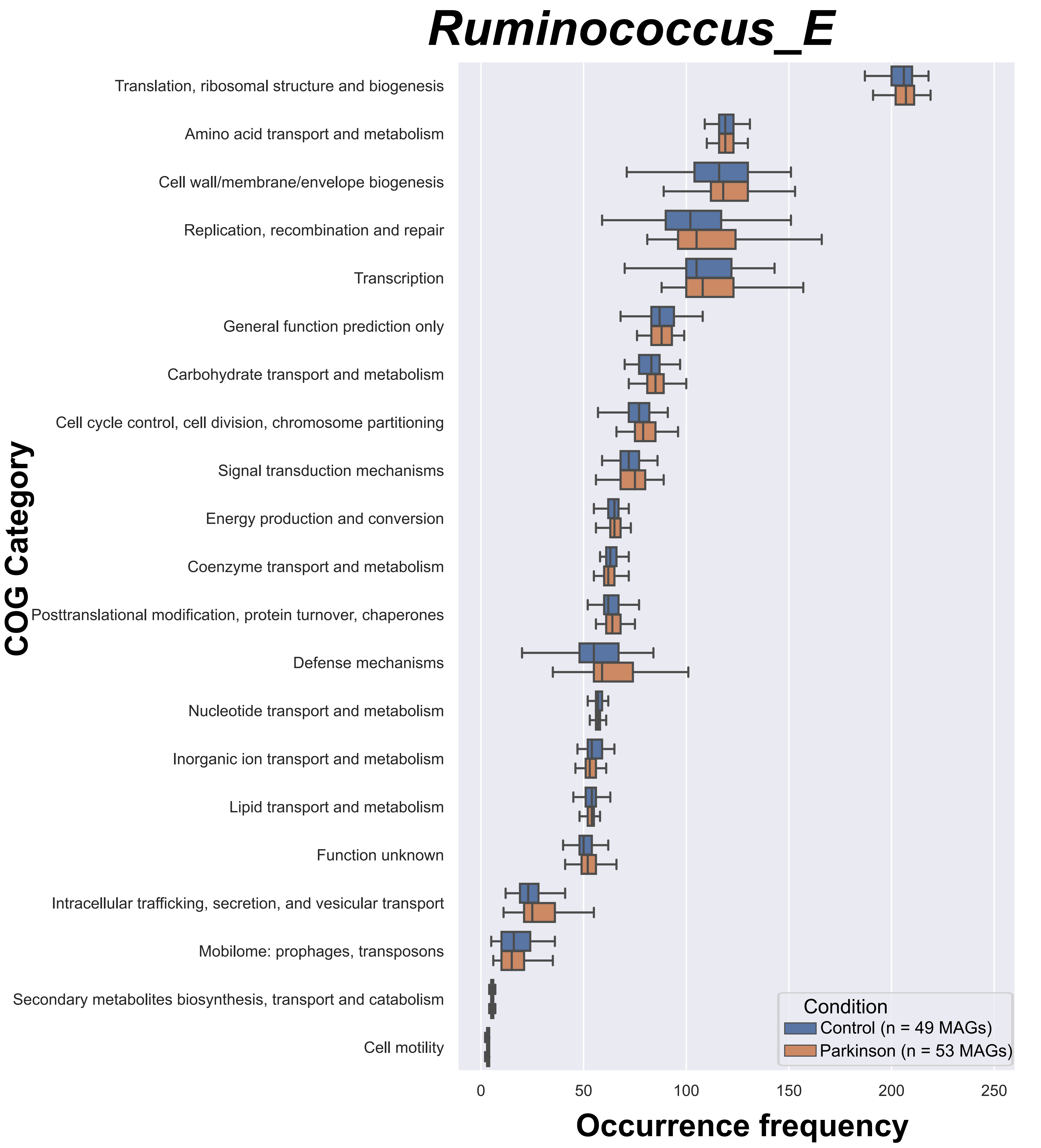

Ruminococcus_E
COG Category
Translation, ribosomal structure and biogenesis
Amino acid transport and metabolism
Cell wall/membrane/envelope biogenesis
Replication, recombination and repair
Transcription
General function prediction only
Carbohydrate transport and metabolism
Cell cycle control, cell division, chromosome partitioning
Signal transduction mechanisms
Energy production and conversion
Coenzyme transport and metabolism
Posttranslational modification, protein turnover, chaperones
Defense mechanisms
Nucleotide transport and metabolism
Inorganic ion transport and metabolism
Lipid transport and metabolism
Function unknown
Intracellular trafficking, secretion, and vesicular transport
Mobilome: prophages, transposons
Secondary metabolites biosynthesis, transport and catabolism
Cell motility
Condition
Control (n = 49 MAGs)
Parkinson (n = 53 MAGs)
0
50
100
150
200
250
Occurrence frequency

Figure S9. The occurrence frequencies of COG categories in selected genera. Each COG category is represented on y axis, and box plot represents the occurrence frequency within the MAGs. Blue is control MAGs, and orange is Parkinson MAGs.
