## Supplementary material for "Metagenome-assembled microbial genomes from Parkinson’s disease fecal samples": All supplementary figures and tables: Supplementary_FigureS12.pdf

a)

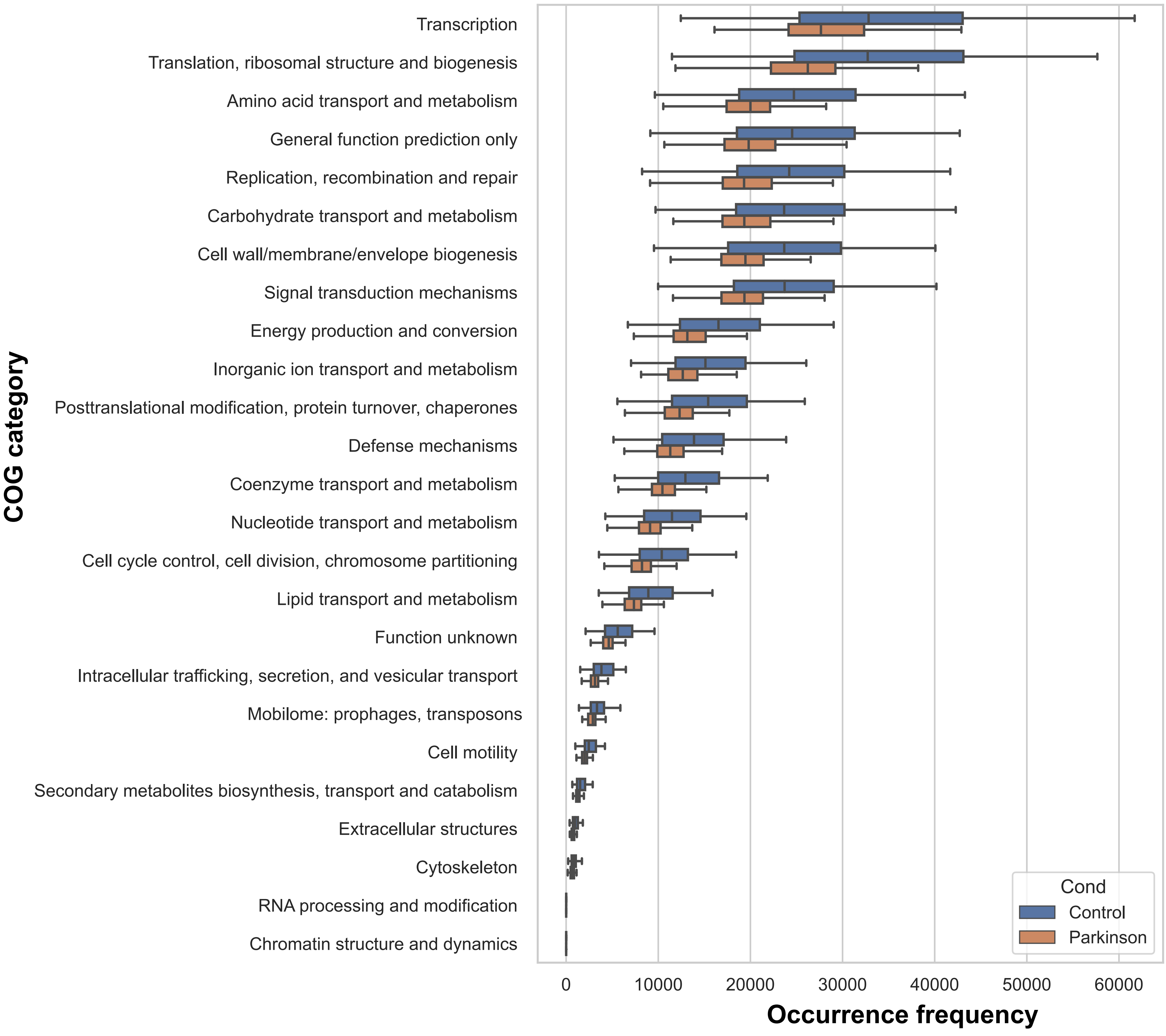

Transcription
Translation, ribosomal structure and biogenesis
Amino acid transport and metabolism
General function prediction only
Replication, recombination and repair
Carbohydrate transport and metabolism
Cell wall/membrane/envelope biogenesis
Signal transduction mechanisms
Energy production and conversion
Inorganic ion transport and metabolism
Posttranslational modification, protein turnover, chaperones
Defense mechanisms
Coenzyme transport and metabolism
Nucleotide transport and metabolism
Cell cycle control, cell division, chromosome partitioning
Lipid transport and metabolism
Function unknown
Intracellular trafficking, secretion, and vesicular transport
Mobilome: prophages, transposons
Cell motility
Secondary metabolites biosynthesis, transport and catabolism
Extracellular structures
Cytoskeleton
RNA processing and modification
Chromatin structure and dynamics
COG category
0
10000
20000
30000
40000
50000
60000
Occurrence frequency
Cond
Control
Parkinson

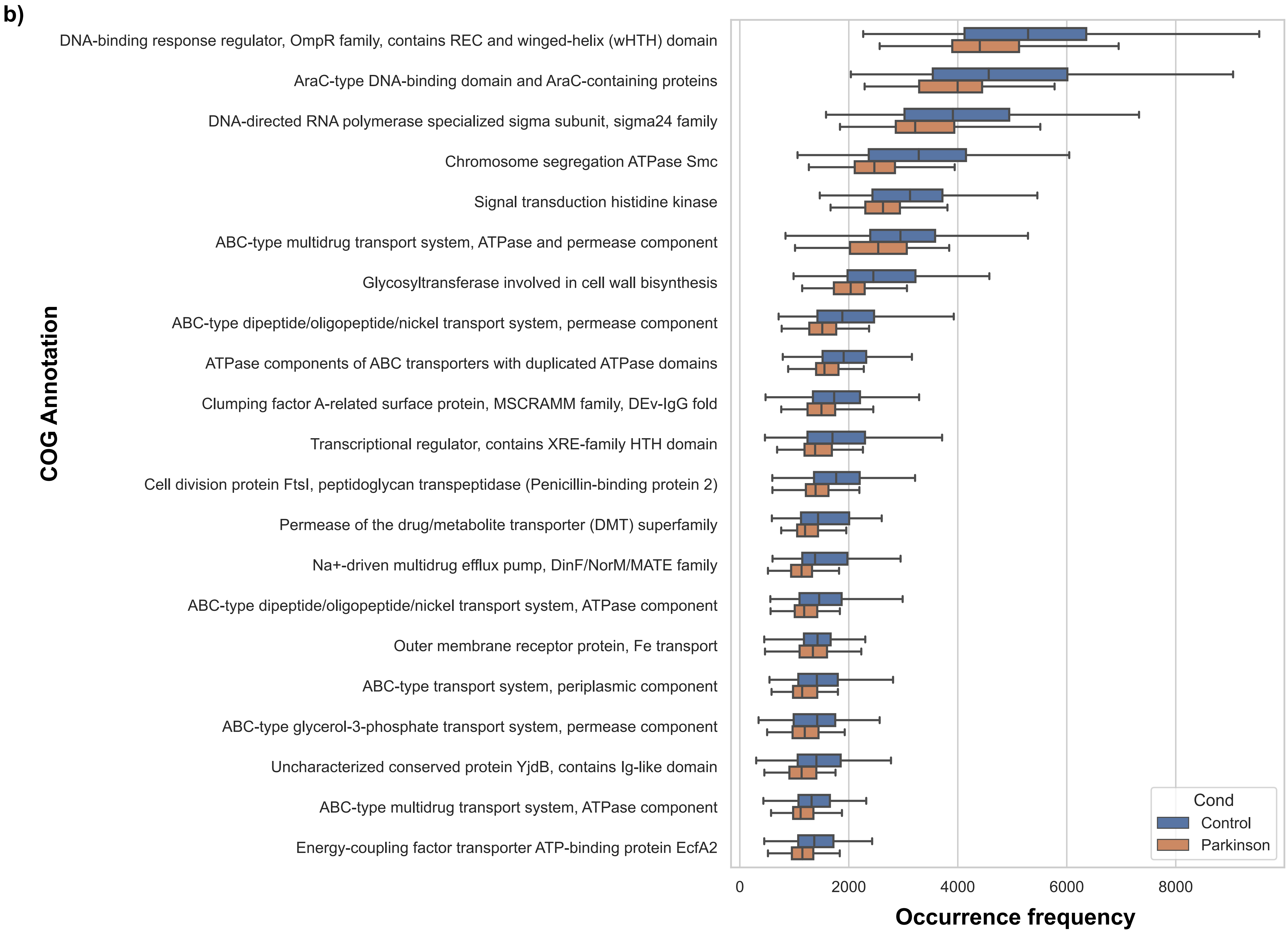
b)
COG Annotation
DNA-binding response regulator, OmpR family, contains REC and winged-helix (wHTH) domain
AraC-type DNA-binding domain and AraC-containing proteins
DNA-directed RNA polymerase specialized sigma subunit, sigma24 family
Chromosome segregation ATPase Smc
Signal transduction histidine kinase
ABC-type multidrug transport system, ATPase and permease component
Glycosyltransferase involved in cell wall bisynthesis
ABC-type dipeptide/oligopeptide/nickel transport system, permease component
ATPase components of ABC transporters with duplicated ATPase domains
Clumping factor A-related surface protein, MSCRAMM family, DEv-IgG fold
Transcriptional regulator, contains XRE-family HTH domain
Cell division protein FtsI, peptidoglycan transpeptidase (Penicillin-binding protein 2)
Permease of the drug/metabolite transporter (DMT) superfamily
Na+-driven multidrug efflux pump, DinF/NorM/MATE family
ABC-type dipeptide/oligopeptide/nickel transport system, ATPase component
Outer membrane receptor protein, Fe transport
ABC-type transport system, periplasmic component
ABC-type glycerol-3-phosphate transport system, permease component
Uncharacterized conserved protein YjdB, contains Ig-like domain
ABC-type multidrug transport system, ATPase component
Energy-coupling factor transporter ATP-binding protein EcfA2
0
2000
4000
6000
8000
Occurrence frequency
Cond
Control
Parkinson

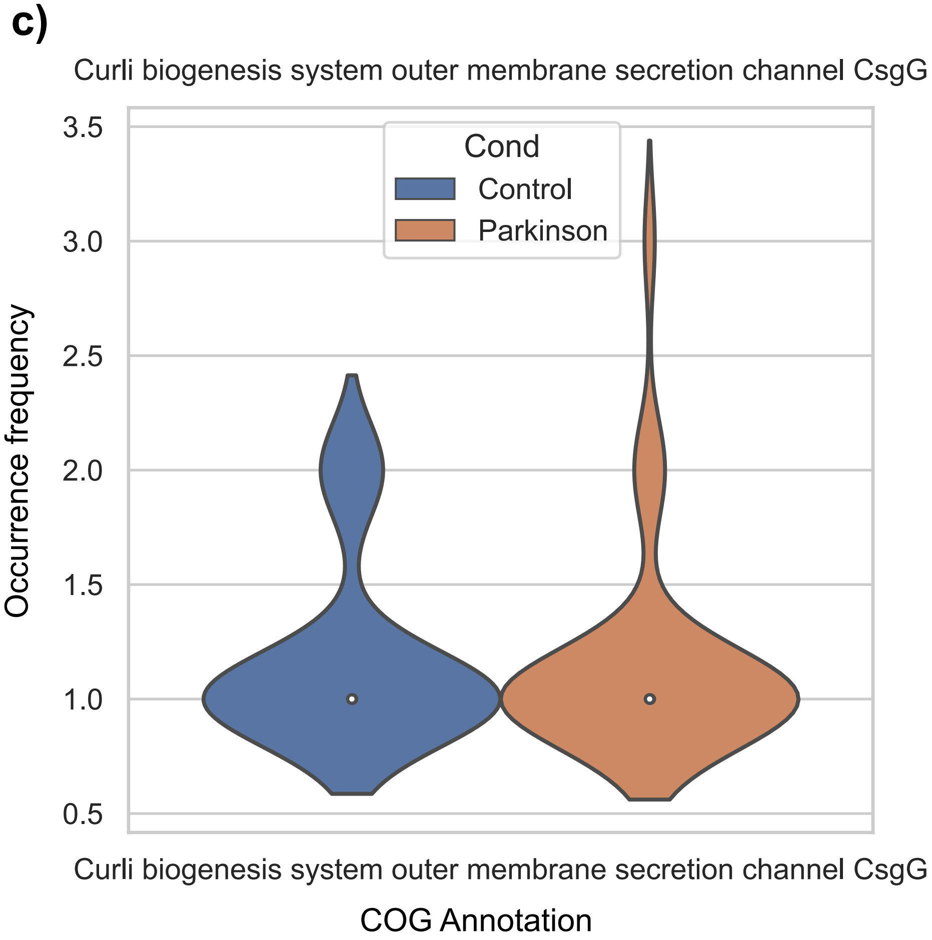
c)
Curli biogenesis system outer membrane secretion channel CsgG
Cond
Control
Parkinson
Occurrence frequency
3.5
3.0
2.5
2.0
1.5
1.0
0.5
Curli biogenesis system outer membrane secretion channel CsgG
COG Annotation

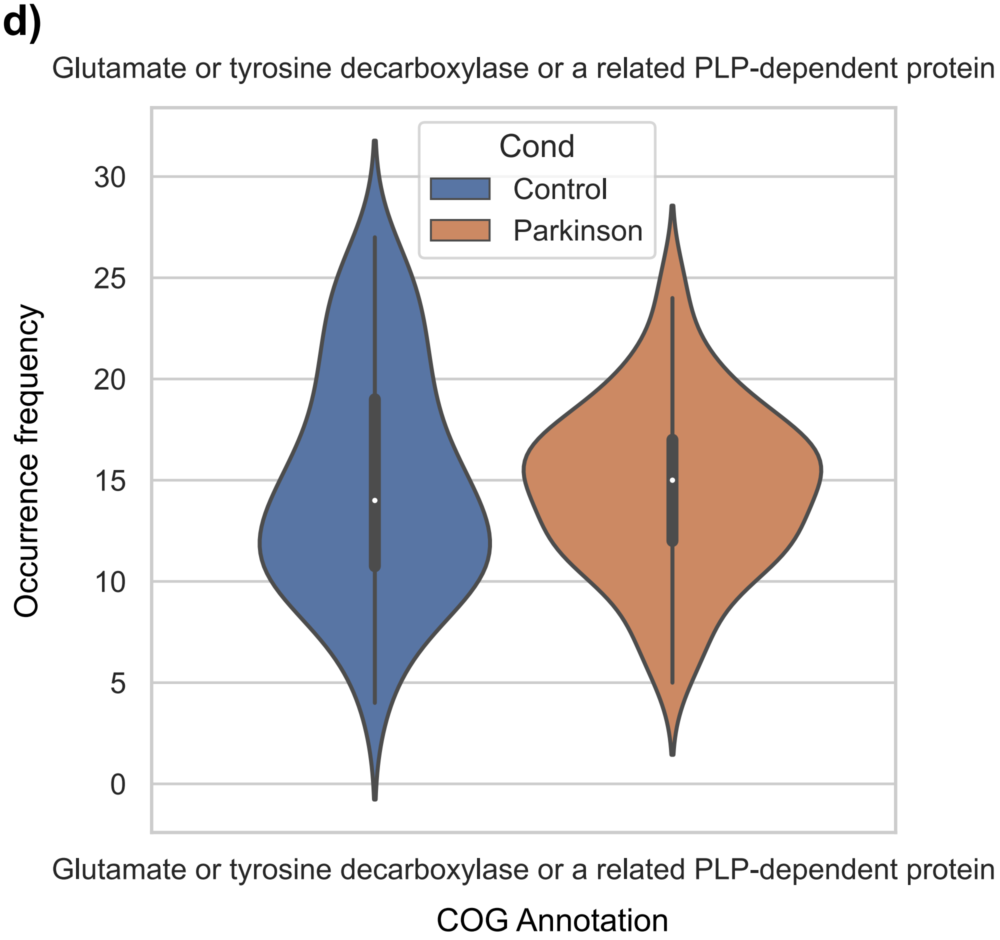
d)
Glutamate or tyrosine decarboxylase or a related PLP-dependent protein
Cond
Control
Parkinson
Occurrence frequency
30
25
20
15
10
5
0
Glutamate or tyrosine decarboxylase or a related PLP-dependent protein
COG Annotation

Figure S12. COG annotation occurrence frequency in all genes within assemblies. a) Top 20 COG category occurrence frequency, b) Top 20 COG annotation occurrence frequency, c) occurrence frequency of “Curli biogenesis system outer membrane secretion channel CsgG”, d) occurrence frequency of “Glutamate or tyrosine decarboxylase or a related PLP-dependent protein”
