## Supplementary material for "Metagenome-assembled microbial genomes from Parkinson’s disease fecal samples": All supplementary figures and tables: Supplementary_FigureS13.pdf

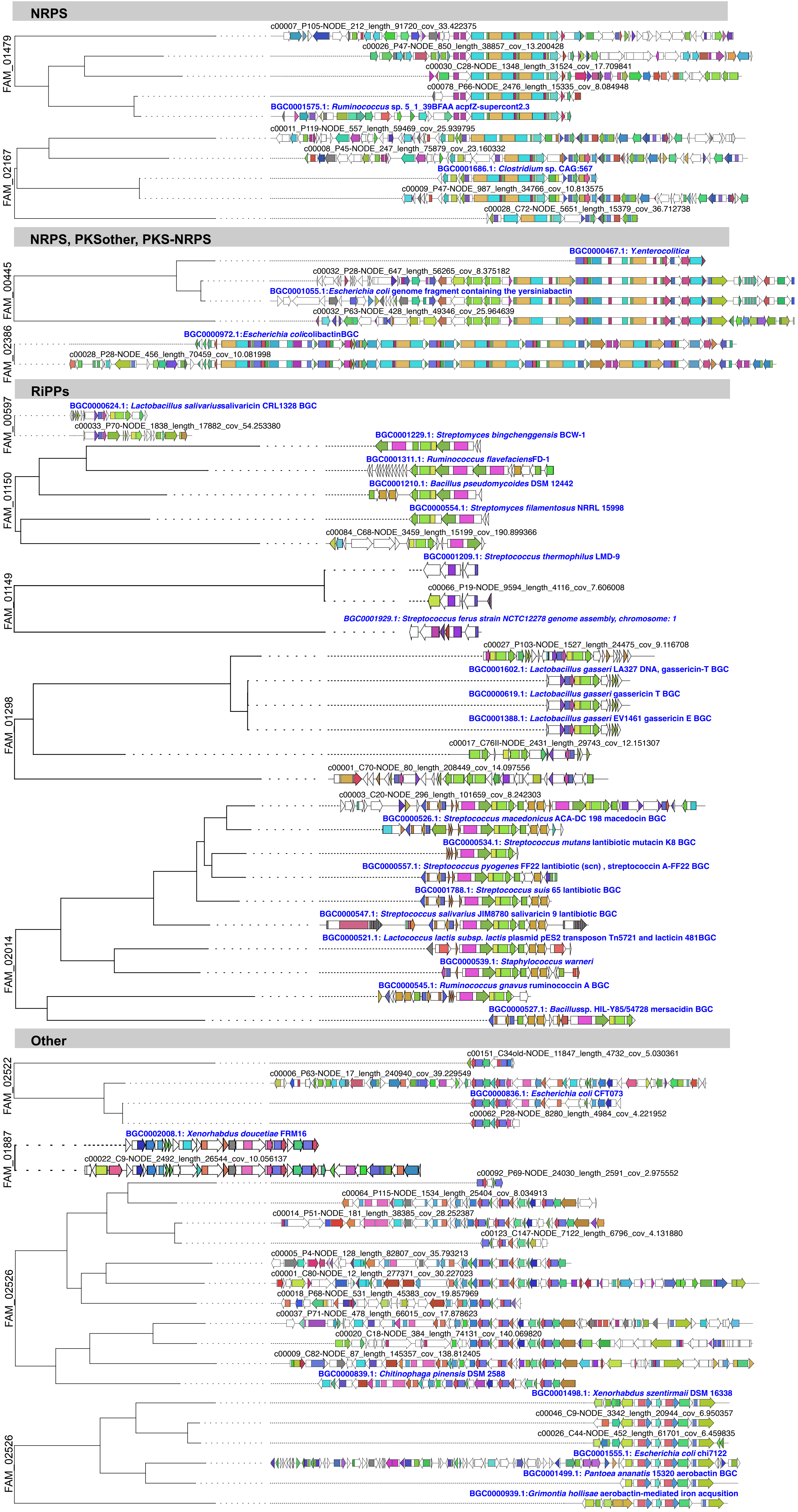

NRPS
FAM_01479
c00007_P105-NODE_212_length_91720_cov_33.422375
c00026_P47-NODE_850_length_38857_cov_13.200428
c00030_C28-NODE_1348_length_31524_cov_17.709841
c00078_P66-NODE_2476_length_15335_cov_8.084948
BGC0001575.1: Ruminococcus sp. 5_1_39BFAA acpfZ-supercont2.3
FAM_02167
c00011_P119-NODE_557_length_59469_cov_25.939795
c00008_P45-NODE_247_length_75879_cov_23.160332
BGC0001686.1: Clostridium sp. CAG:567
c00009_P47-NODE_987_length_34766_cov_10.813575
c00028_C72-NODE_5651_length_15379_cov_36.712738
NRPS, PKSother, PKS-NRPS
FAM_00445
BGC0000467.1: Y.enterocolitica
c00032_P28-NODE_647_length_56265_cov_8.375182
BGC0001055.1:Escherichia coli genome fragment containing the yersiniabactin
c00032_P63-NODE_428_length_49346_cov_25.964639
FAM_02386
BGC0000972.1:Escherichia colicolibactinBGC
c00028_P28-NODE_456_length_70459_cov_10.081998
RiPPs
FAM_00597
BGC0000624.1: Lactobacillus salivariussalivaricin CRL1328 BGC
c00033_P70-NODE_1838_length_17882_cov_54.253380
FAM_01150
BGC0001229.1: Streptomyces bingchenggensis BCW-1
BGC0001311.1: Ruminococcus flavefaciensFD-1
BGC0001210.1: Bacillus pseudomycoides DSM 12442
BGC0000554.1: Streptomyces filamentosus NRRL 15998
c00084_C68-NODE_3459_length_15199_cov_190.899366
FAM_01149
BGC0001209.1: Streptococcus thermophilus LMD-9
c00066_P19-NODE_9594_length_4116_cov_7.606008
BGC0001929.1: Streptococcus ferus strain NCTC12278 genome assembly, chromosome: 1
FAM_01298
c00027_P103-NODE_1527_length_24475_cov_9.116708
BGC0001602.1: Lactobacillus gasseri LA327 DNA, gassericin-T BGC
BGC0000619.1: Lactobacillus gasseri gassericin T BGC
BGC0001388.1: Lactobacillus gasseri EV1461 gassericin E BGC
c00017_C76II-NODE_2431_length_29743_cov_12.151307
c00001_C70-NODE_80_length_208449_cov_14.097556
FAM_02014
c00003_C20-NODE_296_length_101659_cov_8.242303
BGC0000526.1: Streptococcus macedonicus ACA-DC 198 macedocin BGC
BGC0000534.1: Streptococcus mutans lantibiotic mutacin K8 BGC
BGC0000557.1: Streptococcus pyogenes FF22 lantibiotic (scn) , streptococcin A-FF22 BGC
BGC0001788.1: Streptococcus suis 65 lantibiotic BGC
BGC0000547.1: Streptococcus salivarius JIM8780 salivaricin 9 lantibiotic BGC
BGC0000521.1: Lactococcus lactis subsp. lactis plasmid pES2 transposon Tn5721 and lacticin 481BGC
BGC0000539.1: Staphylococcus warneri
BGC0000545.1: Ruminococcus gnavus ruminococcin A BGC
BGC0000527.1: Bacillussp. HIL-Y85/54728 mersacidin BGC
Other
FAM_02522
c00151_C34old-NODE_11847_length_4732_cov_5.030361
c00006_P63-NODE_17_length_240940_cov_39.229549
BGC0000836.1: Escherichia coli CFT073
c00062_P28-NODE_8280_length_4984_cov_4.221952
FAM_01887
BGC0002008.1: Xenorhabdus doucetiae FRM16
c00022_C9-NODE_2492_length_26544_cov_10.056137
FAM_02526
c00092_P69-NODE_24030_length_2591_cov_2.975552
c00064_P115-NODE_1534_length_25404_cov_8.034913
c00014_P51-NODE_181_length_38385_cov_28.252387
c00123_C147-NODE_7122_length_6796_cov_4.131880
c00005_P4-NODE_128_length_82807_cov_35.793213
c00001_C80-NODE_12_length_277371_cov_30.227023
c00018_P68-NODE_531_length_45383_cov_19.857969
c00037_P71-NODE_478_length_66015_cov_17.878623
c00020_C18-NODE_384_length_74131_cov_140.069820
c00009_C82-NODE_87_length_145357_cov_138.812405
BGC0000839.1: Chitinophaga pinensis DSM 2588
FAM_02526
BGC0001498.1: Xenorhabdus szentirmaii DSM 16338
c00046_C9-NODE_3342_length_20944_cov_6.950357
c00026_C44-NODE_452_length_61701_cov_6.459835
BGC0001555.1: Escherichia coli chi7122
BGC0001499.1: Pantoea ananatis 15320 aerobactin BGC
BGC0000939.1:Grimontia hollisae aerobactin-mediated iron acqusition

Figure S13. BGCs from MAGs dereplicated from PD and Control samples that presented similar genes from previously deposited biosynthetic genes from MIBiG.
