## Supplementary material for "Metagenome-assembled microbial genomes from Parkinson’s disease fecal samples": All supplementary figures and tables: Supplementary_Figures_S1_S2_S3_S4_S5.pdf

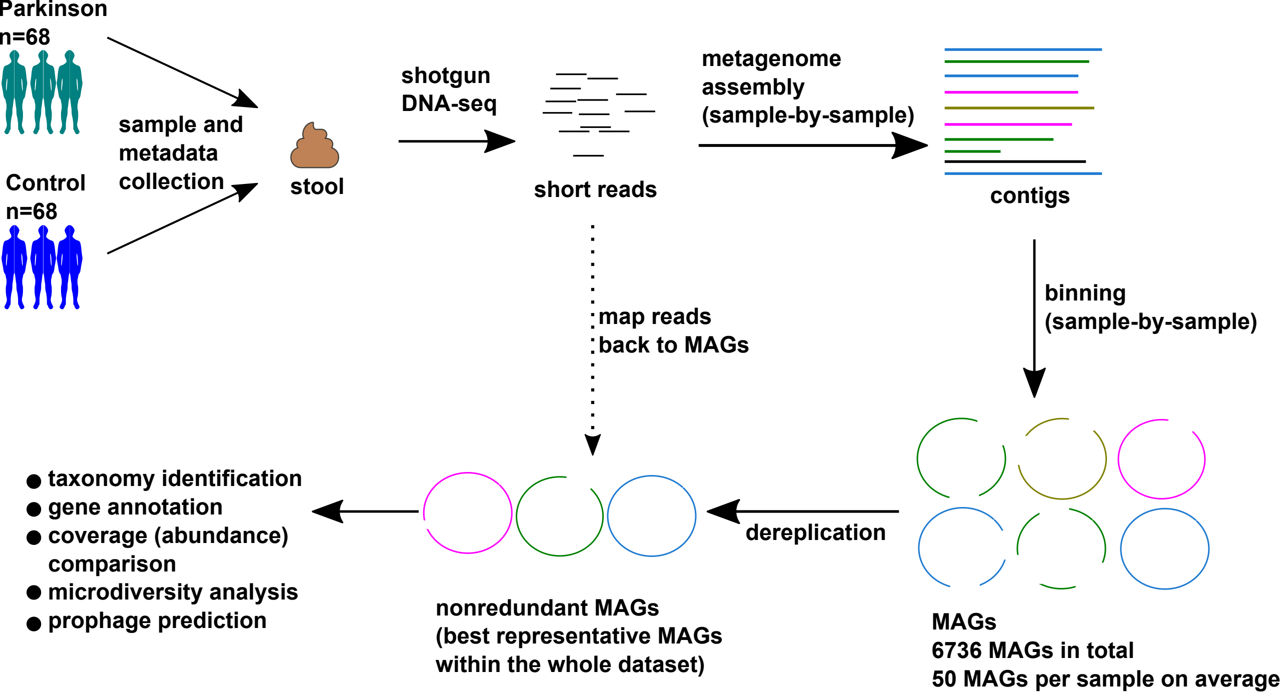


Figure S1. Visual summary of the study.

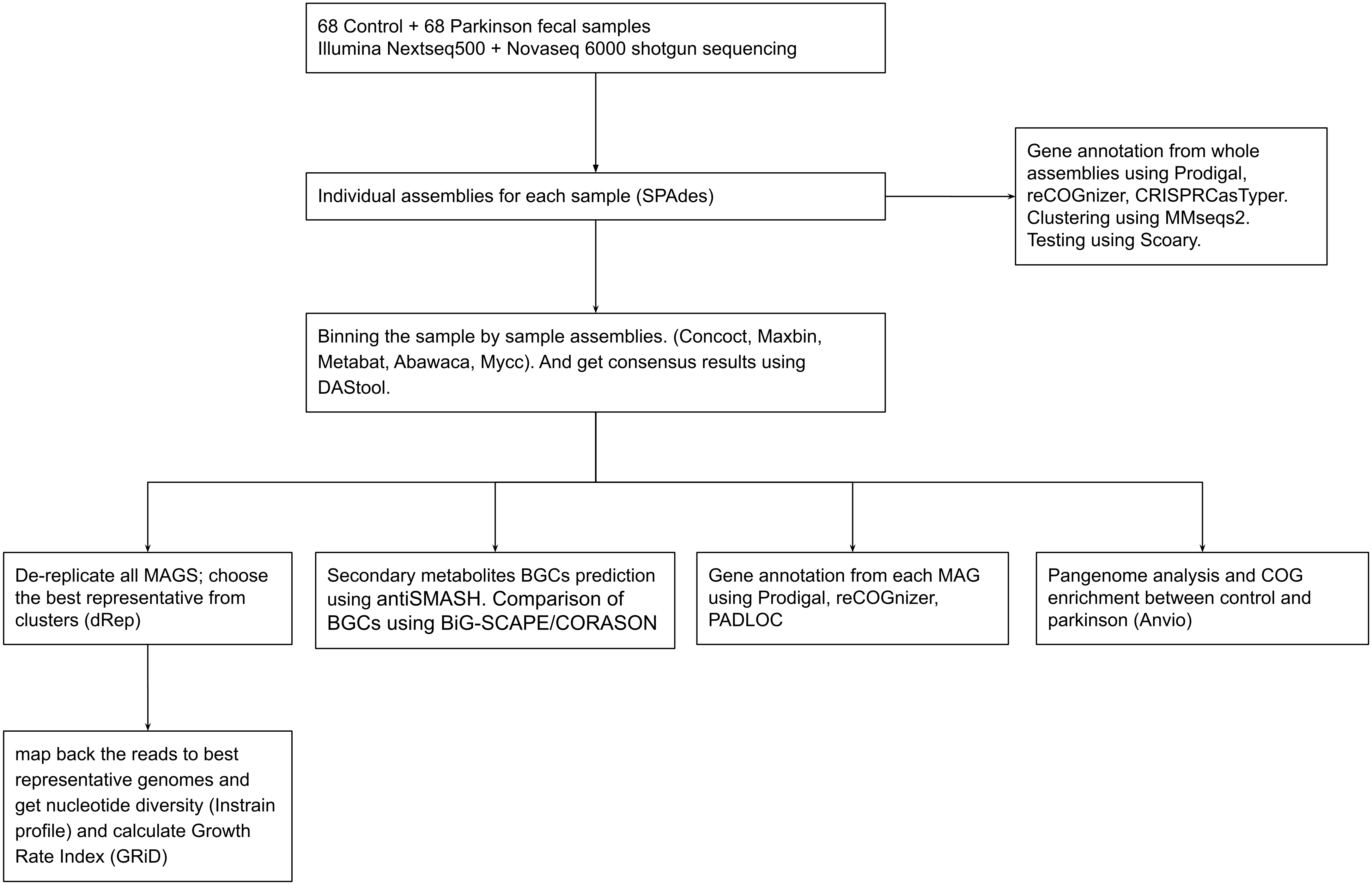


Figure S2. A short summary of the methods that used in this study.

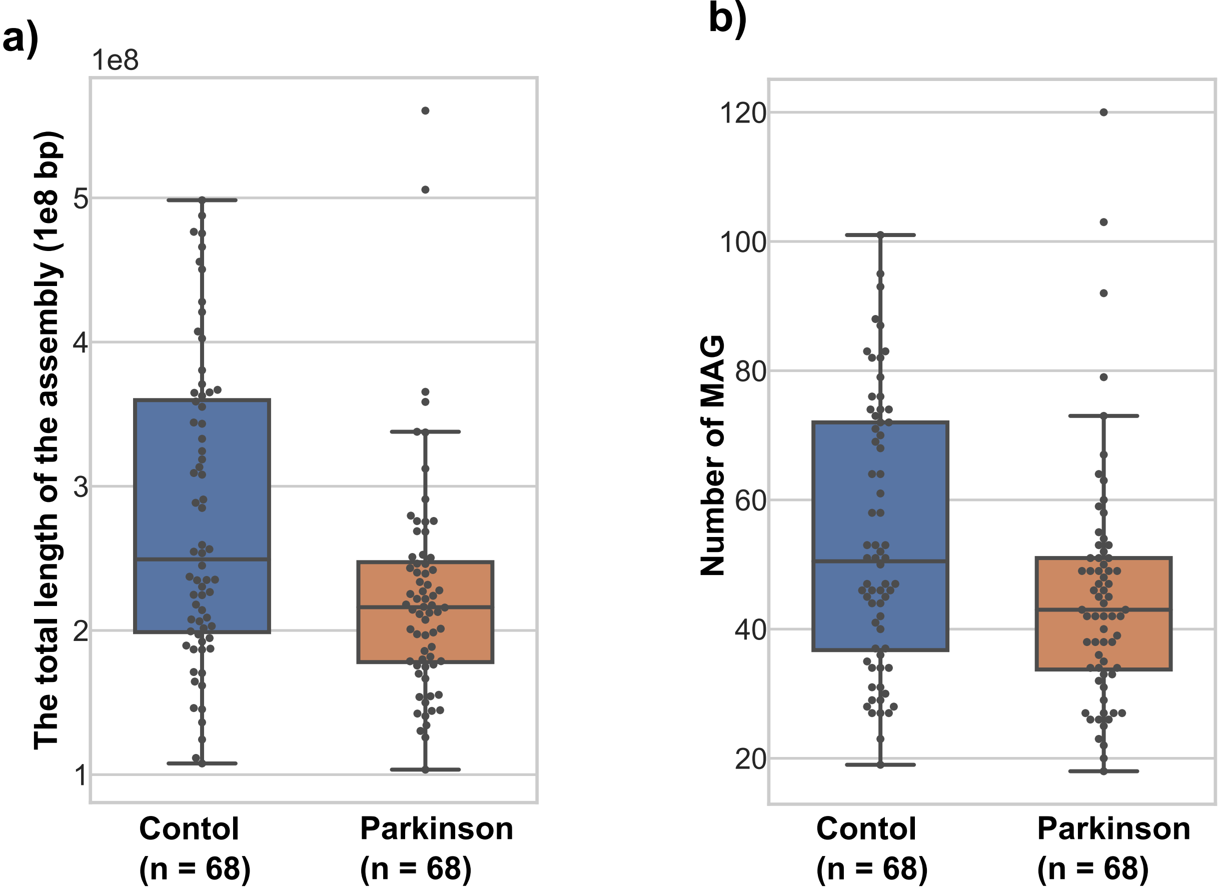


Figure S3. Box-plot showing a) the total length of the assembly, and b) the number of MAGs reconstructed from each sample. Each dot represents one sample. For the total length of the assembly, only contigs larger than 1000 bp are included in the total length calculation.

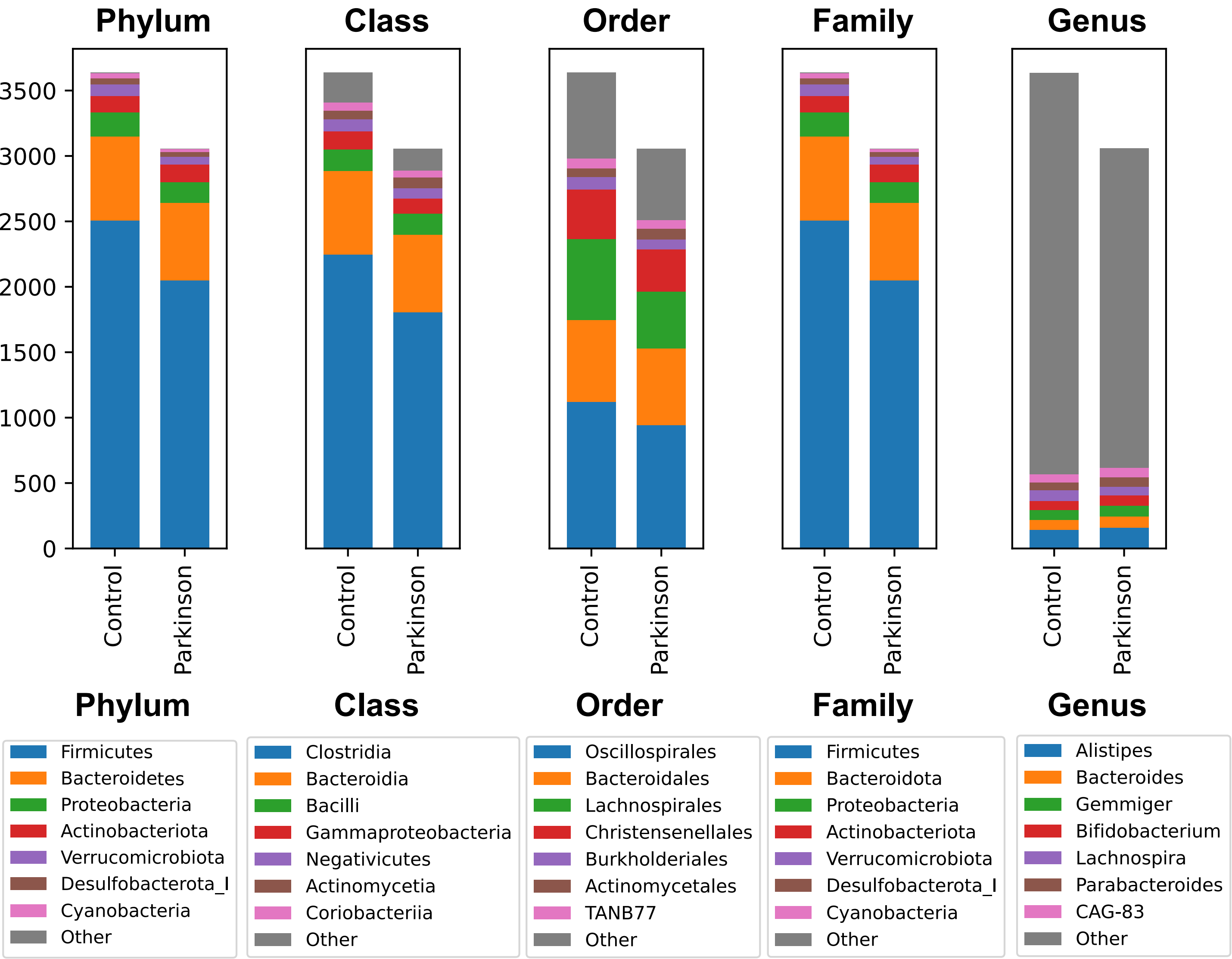


Figure S4. The number of taxonomies of the bacterial MAGs (total of 6692). The bar chart indicates the number of MAGs. The seven most frequently observed taxa are shown in the legend, while the remaining taxonomies were grouped as ‘Other’.

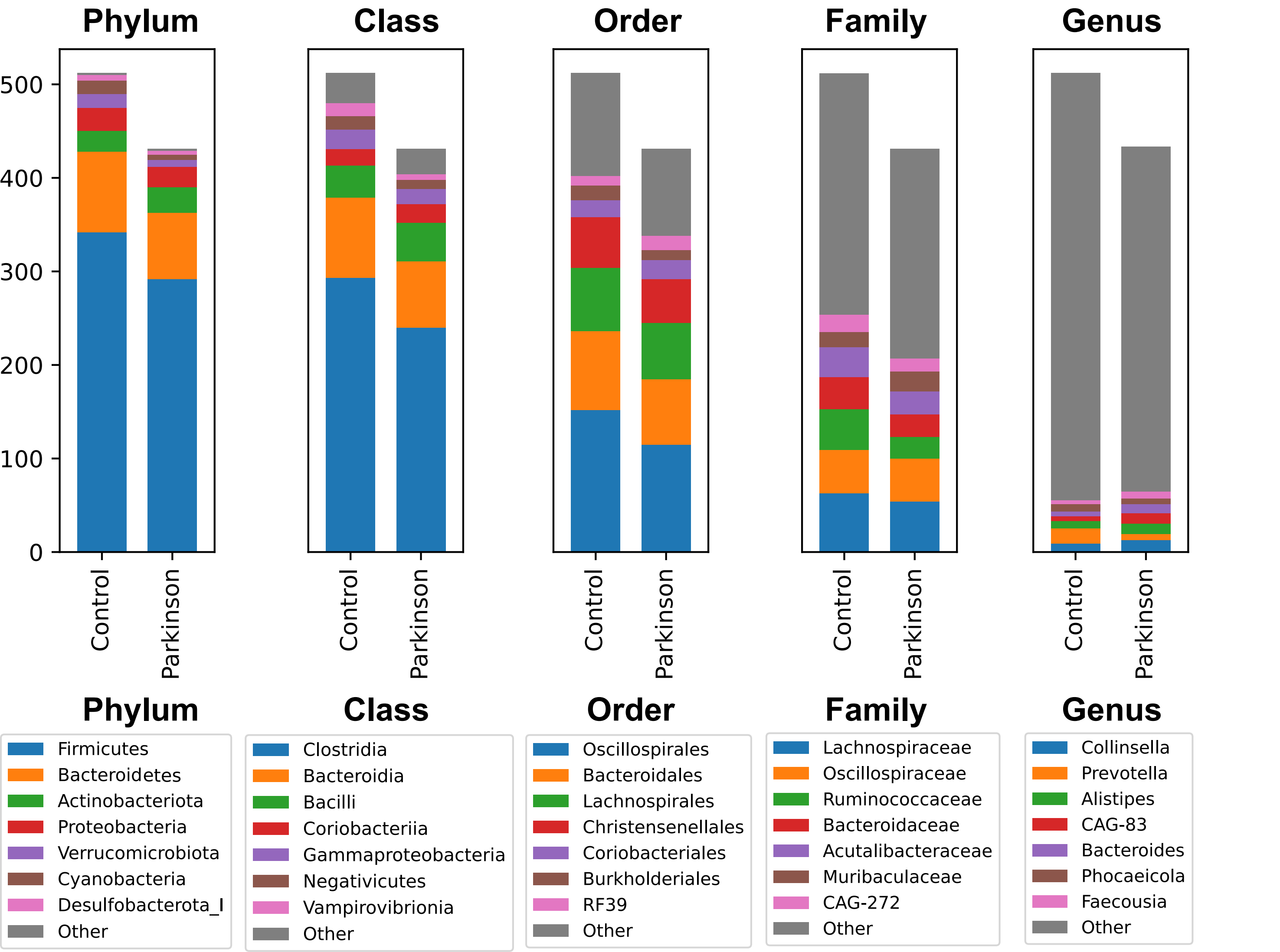


Figure S5. The number of taxonomies of the bacterial dereplicated MAGs (total of 943). The bar chart indicates the number of dereplicated MAGs. The seven most frequently observed taxa are shown in the legend, while the remaining taxonomies were grouped as ‘Other’.
