## Supplementary material for "Metagenome-assembled microbial genomes from Parkinson’s disease fecal samples": All supplementary figures and tables: Supplementary_Figures_S10_S11.pdf

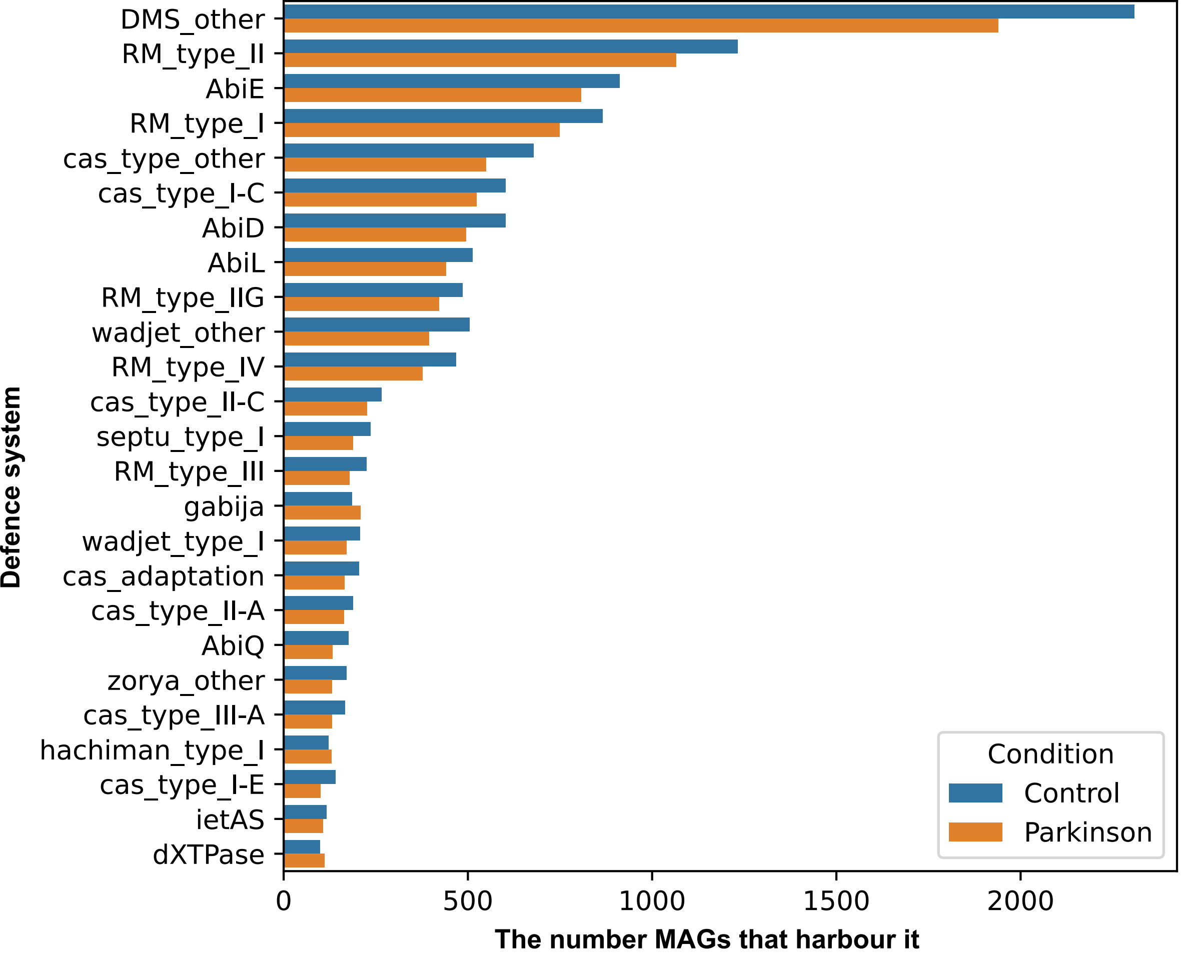


Figure S10. The bar chart shows top 25 defence systems predicted and number of the MAGs that harbour it. Defence system namings are from padlocdb (https://padloc.otago.ac.nz/padloc/systeminformation/). DMS_other represents multiple possible defence systems.

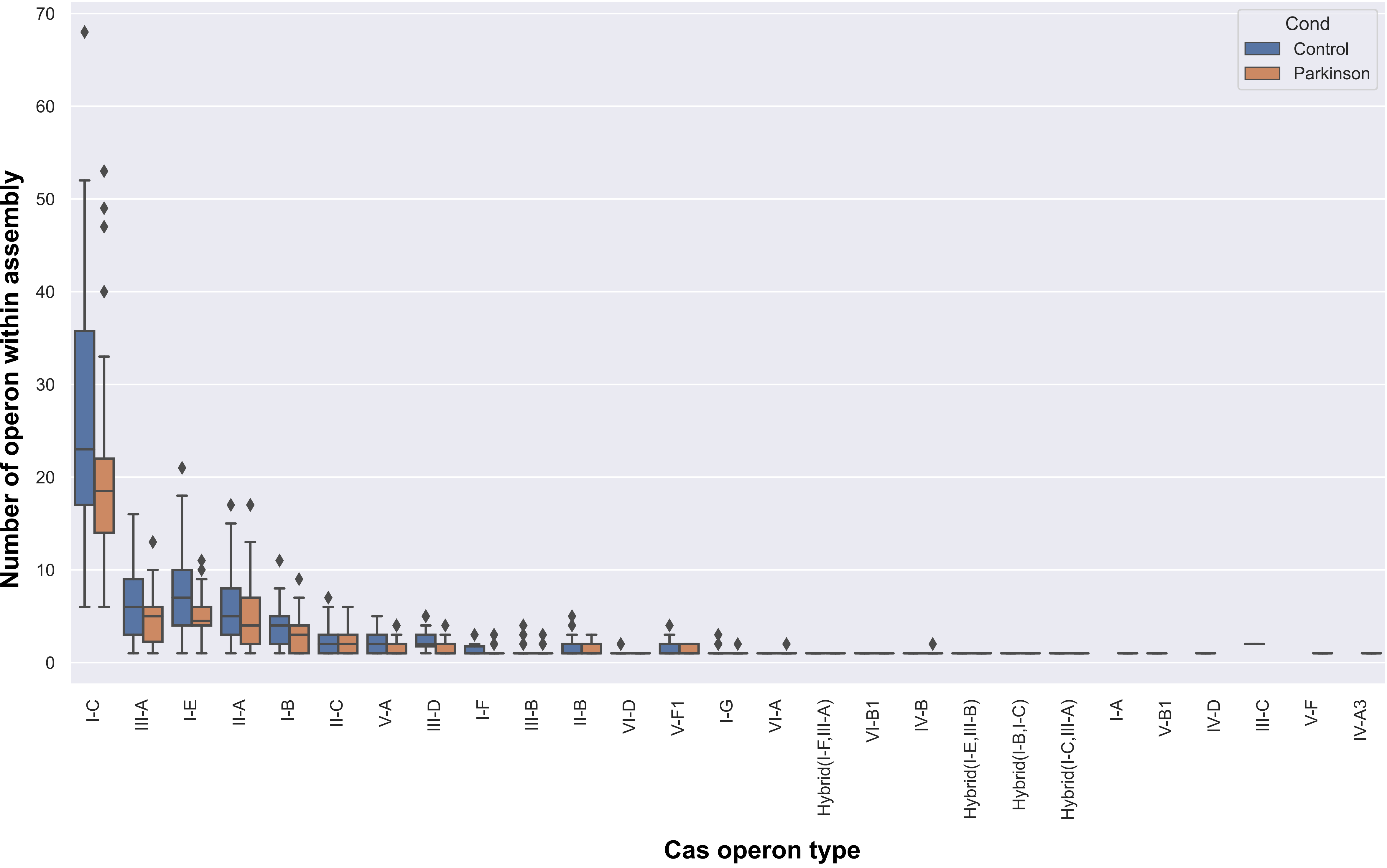


Figure S11.The number of cas operons in each assembly are shown with box plot.
