## Supplementary figures and images for "Metagenome-assembled microbial genomes from Parkinson’s disease fecal samples"

### Supplementary_FigureS14.pdf

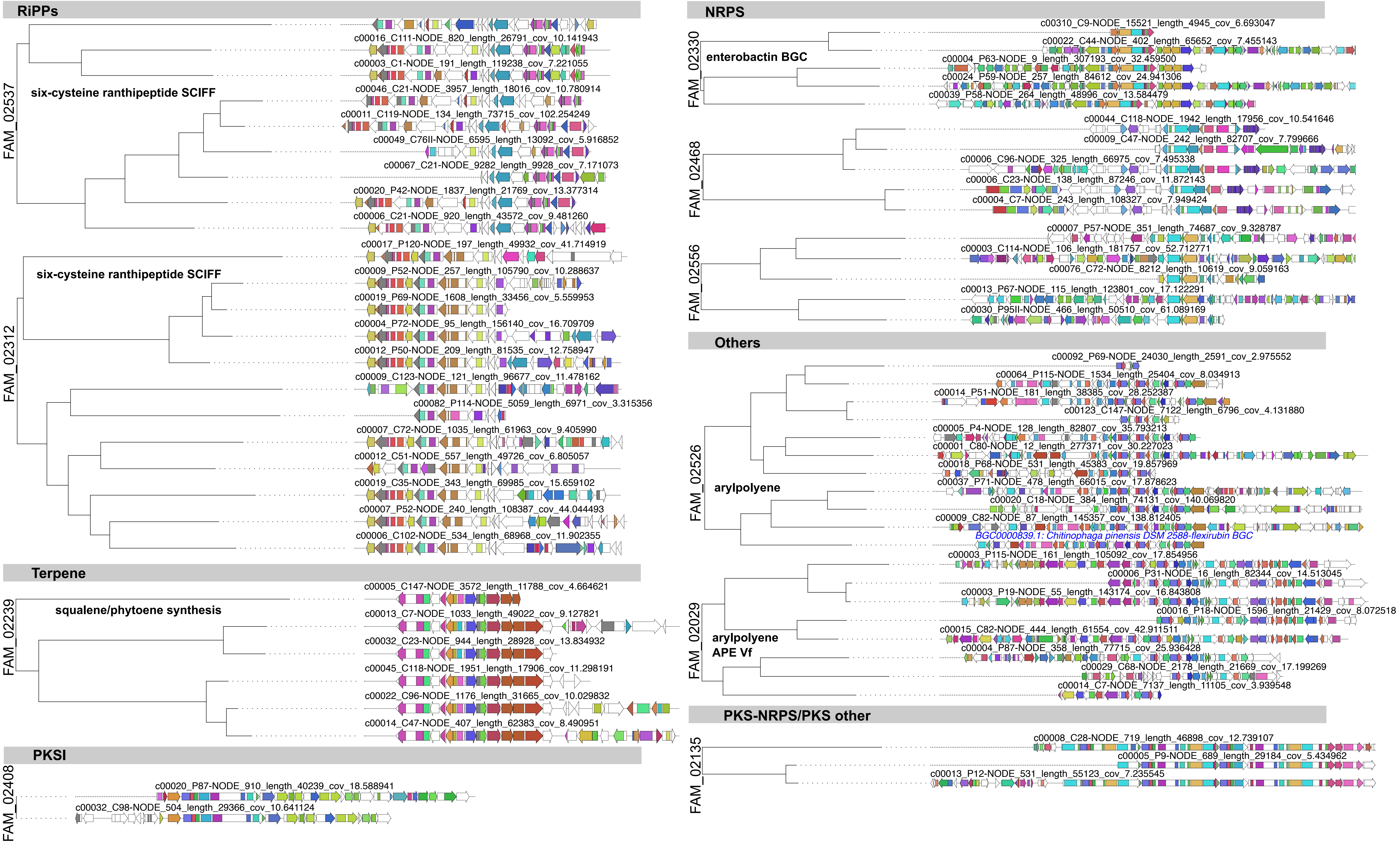

Figure S14. Most commonly secondary metabolites BGCs present in the dereplicated MAGs.
